# Population-scale interpretation of RNA isoform diversity enabled by Isopedia

**DOI:** 10.64898/2026.03.23.713667

**Authors:** Xinchang Zheng, Zev Kronenberg, Sonia Garcia-Ruiz, Ryan M. Layer, Emil K. Gustavsson, Mina Ryten, Fritz J. Sedlazeck

## Abstract

Alternative splicing generates extensive transcriptomic complexity, yet “novelty” is often inflated because of incomplete reference annotations, with 20-70% of transcripts in RNA-Seq studies labeled as novel. Isopedia provides an expandable data structure for reference-agnostic isoform annotation, which we demonstrate here through a population-scale catalog of 1,007 long-read datasets spanning 37 diverse biological contexts. By transitioning from reference-dependent to evidence-weighted annotation, Isopedia provides the frequency-based context necessary to distinguish stochastic noise from biologically active isoforms. In HG002 benchmarks, Isopedia reduced apparent isoform novelty by up to 26-fold, achieving a >95% annotation rate even for low-abundance isoforms typically missed by standard catalogs. The framework further supports systematic exploration of challenging loci such as pseudogenes and gene fusions. Isopedia transforms isoform discovery into a systematic interpretation of the human transcriptome, providing a critical foundation for clinical and functional RNA research. Isopedia is open source and freely available: https://github.com/zhengxinchang/isopedia.

## Introduction

Transcript diversity at each locus drives the functional diversity of multicellular life^1,2^. Through alternative splicing, transcription initiation, and polyadenylation, transcriptional machinery generates multiple isoforms with distinct coding capacities and regulatory roles^3–5^. These isoforms are primary drivers of tissue specialization, developmental timing, and cellular adaptation^6–8^. In human disease, altered isoform usage frequently modifies protein function or regulatory balance in the absence of underlying DNA mutations, making transcript-level variation a critical yet fundamentally under-characterized layer of biological regulation^9–11^.

For decades, short-read RNA sequencing (RNA-seq) was the de facto method for transcriptome profiling^12,13^. A limitation of short-read transcriptomics is the reliance on computational reconstruction of transcripts from fragmented data, resulting in thousands of plausible isoform structures that are challenging to interpret biologically^14–17^. The recent emergence of long-read RNA-seq enables direct sequencing of full-length transcripts from the 5′ to the 3′ untranslated regions (UTRs), resolving complete transcript isoforms and many analytical artifacts of short-read approaches^18,19^. This technological advance has also revealed a challenge: long-read RNA-seq studies consistently report that 26% to 70% of detected isoforms are novel and absent from “gold standard” reference annotations, a pattern also observed, though less pronounced, in short-read transcript assembly^11,20,21^. Such variability suggests that reported novelty rates reflect the incompleteness of reference annotations as much as genuine transcript diversity, exposing a fundamental challenge in how the field defines and interprets isoform discovery^19^.

A fundamental question in RNA-seq experiments is whether a detected isoform is sample-specific or instead recurrent across broader biological contexts, suggesting that it is a functionally relevant transcript. However, current estimates of high isoform novelty likely reflect limited integration of existing transcriptomic evidence rather than true biological novelty, a gap that population-scale indexing of full-length transcripts is uniquely positioned to address. This distinction between novel and recurrent isoform is especially important because isoform-level differences can have substantial functional consequences^2,3,22^. A similar problem was historically faced in human genetics, where the pathogenicity of a single-nucleotide variant could not be interpreted without allele frequencies from population-scale reference catalogs such as gnomAD^23^. While querying individual splice junctions across large-scale short-read repositories has proven valuable^24–26^, these resources cannot resolve full-length isoform structure nor are they easily accessible. In practice, isoform novelty is usually defined relative to reference annotations such as GENCODE or RefSeq^27,28^. However, these resources continue to evolve and differ in curation scope and criteria, resulting in thousands of isoforms present in one but absent from the other^29–31^. This creates a circular and unstable definition of “novelty”: a transcript may be labeled novel because it has not yet been incorporated into a given annotation release, rather than because it represents a rare or context-specific biological event. Despite the growing number of long-read RNA-seq datasets, no population-scale catalog exists to query complete transcript isoforms across studies. Thus, when a candidate isoform is identified in a disease cohort or a specific cell type, there is no easy way to determine if that isoform has been independently observed across the vast set of public transcriptomes.

To resolve these limitations, we present Isopedia, a population-scale framework that indexes long-read alignments of full-length transcriptomes to enable recurrence-based interpretation of RNA isoforms across tissues and studies. Conceptually, Isopedia provides the transcriptomic equivalent of population variant databases, enabling researchers to assess whether candidate isoforms are recurrent across many samples or represent rare, sample-specific events. By indexing disparate datasets into a unified, searchable architecture, Isopedia enables researchers to instantly contextualize their findings against the human transcriptome at large, with its interpretive power growing as additional datasets are contributed by the community. Our scalable and easily extendable approach enables the identification of high-confidence, recurrent isoforms that escape standard annotations and provides the empirical foundation necessary to identify truly novel, sample-specific isoforms. Researchers can query the index directly using GTF files, individual splice junctions, or fusion breakpoints to assess whether a candidate isoform is recurrent or a rare samples-specific event. We demonstrate Isopedia and its utility by indexing a catalog of 1,007 samples generated from long-read RNA datasets. We benchmark Isopedia against state-of-the-art methods, highlighting its computational efficiency and accuracy in isoform discovery and expression analysis.

## Results

### Scalable read-level indexing of the human transcriptome

Isopedia supports three query modes tailored to different biological questions (**Fig. 1a, 1b**). First, the identification of novel and known isoforms. Here isoform queries accept GTF files as input and identify isofrom entries whose splice-junction structures match the queried transcripts within a user-defined wobble tolerance (default 10bp) which accommodates alignment uncertainty at splice boundaries (see **Methods**). Since long-read sequences can still represent fragmented isoforms, Isopedia employs an Expectation-Maximization algorithm to jointly estimate isoform presence and abundance (reported as transcripts per million, TPM) by probabilistically assigning ambiguous reads to their most likely source isoforms (**Fig. 1b**). Second, splice junction queries across the index which accepts a file specifying individual splice junction sites and returns all isofrom entries overlapping those junctions, enabling systematic exploration of junction usage across a population catalog. Third, gene fusion queries support either precise breakpoint coordinates or broader genomic regions, leveraging the supplementary alignment information stored in isofrom entries to identify fusion transcripts. All query modes support parameters to accommodate alignment uncertainty and biological variation at splice boundaries^25,32^ (**Fig. 1b**, **Supplementary Fig. S1**). Each query can be performed across short- or long-read based analysis data.

**Figure 1:**
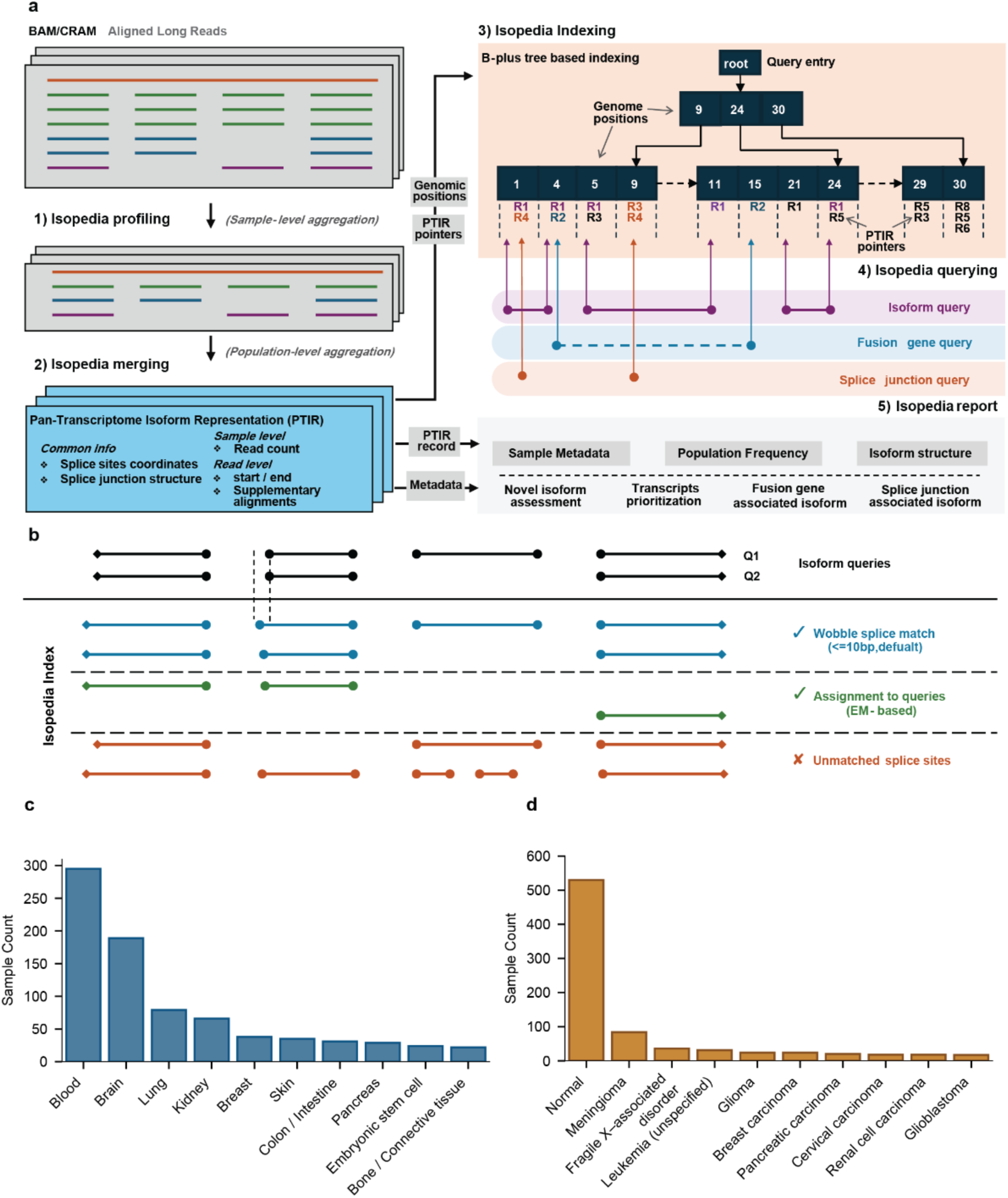
Overview of Isopedia and Isopedia catalog. **a)** The Isopedia workflow. There are three steps in building Isopedia catalogs, (1) extract raw isoform structure from a sample, (2) merge isoform structure across multiple samples to generate a single file archive for all PTIRs, and (3) create a B-plus tree based index on the PTIR genome coordinates and their pointers. The query function has two steps: (4) queried coordinates search and candidate PTIR pointer discovery, and (5) retrieval of matching PNIR records with associated annotations. **b)** Isopedia distinguishes informative long-read alignments from uninformative or noisy reads. Each horizontal line represents a read’s exon segment in reference coordinate space. Circular dots denote splice sites, and diamond-shaped markers indicate read start and end positions. Reads that fully match all splice junctions of a query isoform are uniquely assigned. Reads that partially share splice junctions with the query, without introducing additional junctions, are assigned using an expectation-maximization. Reads containing unmatched or extra splice junctions are excluded from downstream analysis. Black lines indicate the query isoform, blue and green lines represent matched reads, and orange lines indicate unmatched records. **c**) Tissue composition of the 1,007 samples included in the population-scale Isopedia catalog. Top 10 tissues are shown, ordered by their sample count. The labels of tissue were curated and standardized manually. **d**) Disease status distribution of 1,007 samples represented in the Isopedia catalog. Top 10 disease status(including normal tissue) are shown, ordered by their sample count. **PTIR**: pan-transcriptome isoform record; **TPM**: transcripts per million.

To enable population-scale analyses of isoforms, we aggregated 1,007 public human long-read RNA sequencing datasets (603 PacBio, 404 ONT), manually curated and normalized their metadata, and generated a structured, community-available catalog. This catalog spans 37 tissues, with blood (295 samples), brain (189), lung (79), and kidney (66) being the most represented (**Fig. 1c**). The majority of samples were derived from healthy individuals (530), with meningioma (84) being the most common disease condition (**Fig. 1d**). Samples were obtained from ENCODE^21^ (107) and SRA^33^ (900, deposited 2020-2025), and metadata including tissue type and disease status were manually standardized (**Supplementary Table S1**). This catalog serves as the reference for all population-level analyses presented herein and is publicly available: https://github.com/zhengxinchang/isopedia.

### Isopedia achieves robust and efficient isoform re-identification and quantification

We validated the Isopedia query functionality with the in silico long-read benchmark from LRGASP^19^. We first assessed if queries returned the correct number of isoforms and not false positives (i.e. sensitivity and specificity, see **Methods**). Isopedia achieved the highest F1 score (0.981) for PacBio data compared with other methods (Oarfish^34^:0.980, StringTie^16^:0.943, IsoQuant^35^:0.947, Bambu^36^:0.932). For the ONT data benchmark, Oarfish had the highest F1 score (0.962), followed by Isopedia (0.894), StringTie (0.876), and IsoQuant (0.688) (**Fig. 2a, Supplementary Table S2**). The simulated ONT reads had shorter median lengths and a higher proportion of truncated alignments compared to PacBio, and Oarfish, unlike the other tools, aligned reads directly to the ground-truth transcripts, providing an uncorrected advantage in this evaluation. LRGASP also included a real-world dataset with technical replicates, which we used to assess Isopedia’s query accuracy by evaluating concordance across replicates. We observed an average consistency rate of 91.5% across the 18 replicates, equivalent to other tools (**Supplementary Note 1**, **Supplementary Table S3**).

**Figure 2:**
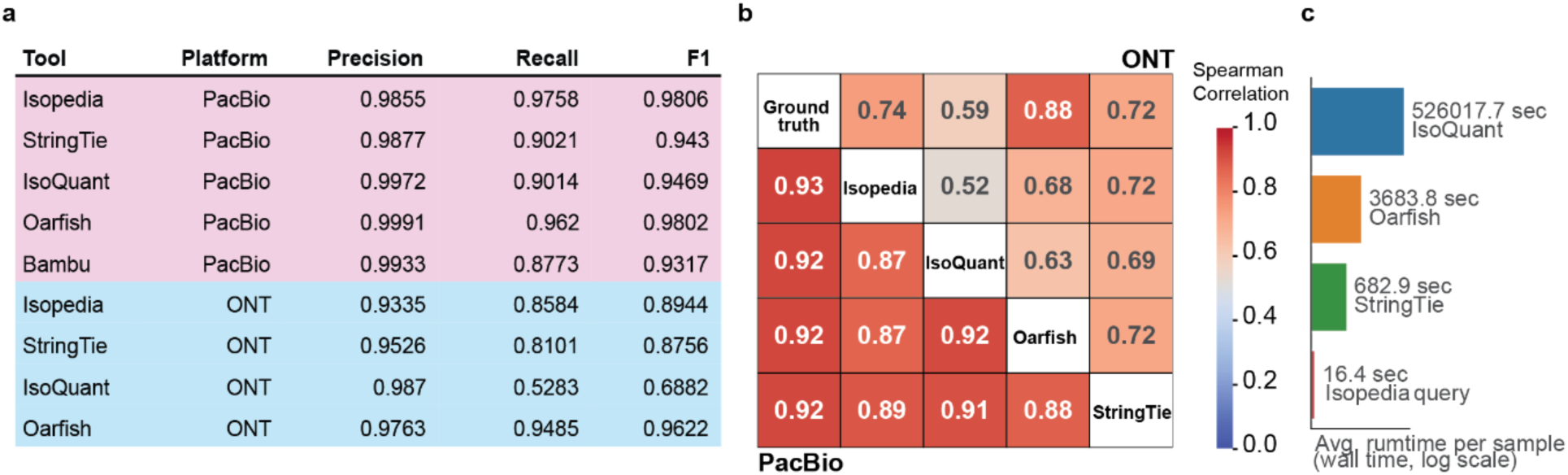
Query performance assessment of Isopedia. **a)** Isopedia performance on LRGASP simulated PacBio and ONT datasets. **b)** Spearman correlation of transcript abundances between Isopedia, ground truth, and other tools. The lower left triangle represents PacBio data; the upper right triangle represents ONT data. **c)** Computational resource comparison of tools. Average runtime (wall time) in seconds was measured.

Quantification accuracy was also validated using the LRGASP ground truth set. We compared the TPM value across all tools against the ground truth using Spearman correlation coefficient. For PacBio data (**Fig. 2b)** all tools achieved similar performance (Isopedia: r = 0.93; Oarfish, StringTie, IsoQuant: r = 0.92). For ONT, we observed a weaker correlation with ground truth; Isopedia achieved a correlation of 0.74, ranking second after Oarfish (**Fig. 2b**). On PacBio data, Isopedia showed highest correlation with StringTie (r = 0.89), followed by IsoQuant (r = 0.87) and Oarfish (r = 0.87). For ONT datasets, correlations with Isopedia were lower: StringTie (r = 0.72), Oarfish (r = 0.68), and IsoQuant (r = 0.56). StringTie showed the highest correlation with Isopedia for both PacBio and ONT (Fig. 2b). Lastly, we also assessed the quantification across the 18 replicates from the cell line from LRGASP data. We observed an average coefficient of variation of 0.21 across all library types, comparable to that of other tools (**Supplementary Note 1**). Across all query modes, Isopedia was between 42X and 32,00X faster than other methods (**Fig. 2c, Supplementary Table S12**). Isopedia showed comparable accuracy to commonly used RNA-seq analysis tools on individual experiments, enabling its application to population-scale interpretation across large sample collections.

### Isopedia reduces annotation-dependent isoform novelty

A common issue with long-read based RNA analysis is the high fraction (up to 26-70%) of novel isoforms being reported in each paper, which poses a substantial challenge for interpretation in the absence of the population-scale context^11,20,21^. To assess how population-scale context affects isoform novelty estimates, we compared annotation outcomes for HG002 RNA-seq data using traditional reference-based approaches (GFFCompare^37^ against GENCODE V49 and RefSeq Release 110) versus Isopedia’s 1,007-sample catalog. We processed transcripts from HG002 sequenced with PacBio, ONT, and Illumina from the Genome in a Bottle (GIAB) consortium. Transcripts were assembled using StringTie with GENCODE V49 as the reference annotation, yielding 33,920 transcripts from PacBio, 25,061 from ONT, and 31,426 from Illumina. There was low consensus among the sequencing technologies (**Fig. 3a**), with only 8,010 transcripts shared across all technologies and 8,767 transcripts supported by at least two technologies. We evaluated isoform annotation performance across sequencing technologies. Using GFFCompare with GENCODE (V49), we found that 26.7%, 45.54%, and 45.54% of isoforms were classified as novel for PacBio, ONT, and Illumina, respectively; with RefSeq (Release 110), the proportions were 42.86%, 51.55%, and 50.51% for PacBio, ONT and Illumina, respectively. Isopedia identified far fewer potential novel isoforms: only 1.98%, 2.46%, and 4.84% for PacBio, ONT, and Illumina, respectively (**Fig. 3b**). Thus, Isopedia reduced the rate of novel isoform predictions by up to 26-fold compared to conventional approaches (GENCODE: 9.41-13.48 fold, RefSeq: 10.44-26.04 fold).

**Figure 3:**
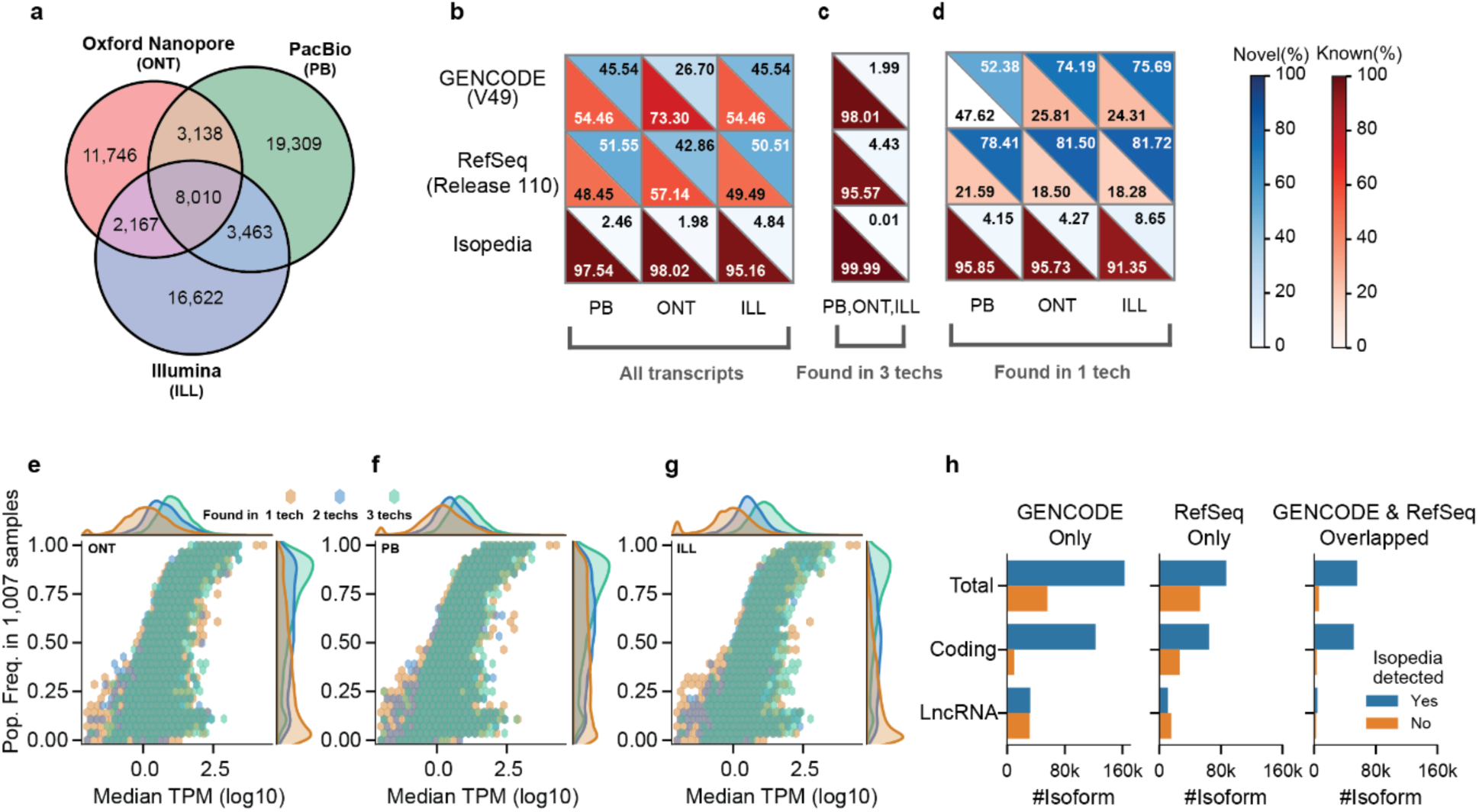
Population-scale re-annotation of HG002 transcripts and reference isoforms using Isopedia. **a)** Overlap of HG002 isoforms detected from HG002 samples that sequenced by ONT, PacBio, and Illumina technology. **b)** Classification of HG002 transcripts by annotation status across all transcripts identified by each sequencing technology (PB, ONT, ILL). Annotation was performed using GENCODE V49 (first row), RefSeq Release 110 (second row), and Isopedia (third row). In each cell, the upper triangle shows the percentage of transcripts classified as novel, and the lower triangle shows the percentage classified as known. Color legends are shared across panels b-d. **c)** As in b), but restricted to transcripts detected by all three technologies. **d)** As in b), but restricted to transcripts uniquely detected by a single technology. **e)** Isoform expression levels (median TPM across all samples) and population frequency from Isopedia for PacBio-derived transcripts. Each transcript is colored according to the number of supporting technologies (1 tech: yellow, 2 techs: blue, 3 techs: green). The top density plot depicts the distribution of expression levels and the right density plot shows the population frequency distribution, both stratified by technology support. Color legends are shared across panels e-g. **f)** As in e), but for ONT-derived transcripts. **g**) As in e), but for Illumina-derived transcripts. **h)** Re-identification of GENCODE only (left), Overlapped between GENCODE and RefSeq (middle) and RefSeq only (right) isoforms from the Isopedia catalog. Total transcripts, protein coding, and lncRNA transcripts are shown. **PB**: PacBio; **ONT**: Oxford Nanopore Technology; **ILL**: Illumina.

Interestingly, there was a high degree of variability in terms of isoform detection rate across the three technologies when using the traditional methods to analyze HG002 (**Fig. 3a**). Nonetheless, we observed strong concordances between reference annotations and Isopedia, as shown in **Fig. 3c**, with all of them achieving annotation rates exceeding 95%. With further investigation, we noted that isoforms only found in one technology against GENCODE or RefSeq, had a very low detection rate of 18-47% (**Fig. 3d**). In contrast, a moderately higher detection rate (74-98%) was observed for isoforms supported by two or more technologies (**Supplementary Fig. S10**). Isopedia showed consistently high detection rates for single technology support (91-95%) and across isoforms reported by two sequencing technologies (99%). Next, we investigated the population frequency and expression of isoforms supported by a single technology. We found that such isoforms tended to be expressed at lower levels and had a lower population frequency in our catalog (**Fig. 3e, 3f,** and **3g**). However, population frequency and expression level are not interchangeable. A lowly-expressed isoform can nonetheless be consistently detected across individuals. This underscores population frequency, as captured by Isopedia, as an independent measure for evaluating isoform reliability. Together, these results demonstrate that Isopedia provides stable, technology-agnostic detection across isoforms of varying abundance and population frequency, particularly where reference-based approaches show reduced sensitivity.

Rather than treating annotation choice as ground truth, Isopedia provides population-scale long-read evidence to quantify which transcript models are reproducibly observed, enabling evidence-weighted interpretation of ‘known’ and ‘novel’ isoforms. Nevertheless, established reference genome annotations are not replaceable and are highly important. Thus, we evaluated the support of GENCODE and RefSeq by their overlap and re-identification rates based on the Isopedia catalog across 1,007 long-read RNA datasets. Furthermore, we stratified isoform support by the population frequency. The GENCODE and RefSeq GTF files contain 280,000 and 201,189 transcript records, respectively, and are known to be overlapping and disagreeing^30,38,39^. We found 61,458 transcripts to be identical between the two resources (class code “=”, GFFCompare), representing 21.95% of GENCODE and 30.55% of RefSeq annotations. Overall, we found 77.92% (218,167) of GENCODE-derived transcripts and 70.88% (142,601) of RefSeq-derived transcripts in the Isopedia catalog.

To better understand how annotation support varies across resources, we stratified transcripts by annotation concordance and transcript type (including protein-coding and lncRNA). Transcripts annotated in both resources showed the highest level of support in the Isopedia catalog with Isopedia supporting 94.02% of these shared protein-coding transcripts (**Fig. 3h**, **Supplementary Table S4** and **S5**). Transcripts unique to a single annotation resource still showed high support in Isopedia, with 92.70% of GENCODE-only and 71.23% of RefSeq-only protein-coding transcripts supported. Of the 61,833 (22.08%) GENCODE transcripts not supported by the Isopedia catalog, the majority were classified as manually annotated (level 2; 80.04%), followed by automatically annotated (level 3; 11.31%) and validated (level 1; 8.65%). Similarly, of the 58,588 (29.12%) unsupported RefSeq transcripts, the majority belonged to the model class (60.96%) rather than the known class (39.04%). In terms of biotype composition, unsupported GENCODE transcripts were predominantly lncRNAs (54.50%) and protein-coding genes (21.05%), whereas unsupported RefSeq transcripts were predominantly protein-coding genes (49.48%) and lncRNAs (31.04%).

Given the relatively low annotation rate (71.23%) observed for RefSeq protein-coding transcripts, we further examined 115,588 transcripts according to their evidence level in RefSeq. Isopedia identified 62,754 (82.90%) manually curated transcripts and 52,821 (76.70%) automatically annotated transcripts. Curated transcripts were significantly (Fisher’s Exact Test, *P* < 0.001) enriched among Isopedia-supported transcripts, showing that well-curated annotations are more likely to be represented in the Isopedia catalog. This was further supported by the analysis of MANE (Matched Annotation from NCBI and EMBL-EBI) transcripts^31^ (GENCODE:18,092, RefSeq:18,209), which revealed that the vast majority (GENCODE: 92.83% and RefSeq: 92.53%) of these clinically curated models were supported by Isopedia, highlighting its strong concordance with high-confidence reference annotations while extending beyond them to capture additional isoform diversity. Overall, we found that GENCODE had the most long-read support in the Isopedia catalog.

### Population-scale indexing reveals constrained and relaxed isoform diversity

To illustrate how population-scale indexing enables biological discovery beyond isoform annotation, we used Isopedia to explore genome-wide patterns of isoform diversity across genes and pseudogenes. We hypothesized that a lack of selective pressure would be expected to generate higher pseudogene isoform diversity compared to constrained protein coding equivalents. We first validated this hypothesis on a well-known medically relevant gene *GBA1* and its paired pseudogene *GBAP1*. Previous studies have revealed unappreciated *GBA1/GBAP1* isoform diversity by directly counting isoforms from long-read data^40^. Gustavsson et al. reported this pattern over *GBA1* and *GBAP1* where they identified 32 and 48 isoforms for *GBA1* and *GBAP1* respectively, across 29 samples (**Fig. 4a**)^40^. In contrast, GENCODE and RefSeq show the opposite for *GBA1* vs. *GBAP1* of lower isoforms count of the pseudo gene compared to the coding gene (GENCODE: *GBA1*:17, *GBAP1*:10; RefSeq: *GBA1*:12, *GBAP1*:2) (**Fig. 4a**). We reanalyzed all candidate transcripts (*GBA1*: 2,368 candidates, *GBAP1*: 3,083 candidates) from Gustavsson et al. by querying them against the Isopedia catalog. Based on evidence from full-splice matches, we re-identified 86 *GBA1* and 303 *GBAP1* candidate isoforms in at least two independent Isopedia catalog samples (**Fig. 4a**; see **Supplementary Note 2, Supplementary Table S13** and **S14** for details), suggesting that the isoform numbers at these loci are substantially greater than previously appreciated. While the above isoform-level query validates known candidates, it is inherently limited to previously reported transcripts. To minimize annotation-dependent bias in estimating isoform diversity, we used Isopedia’s splice query function to systematically enumerate isoforms supported in at least two independent samples (mapping quality >5) and from both sequencing technologies. Isoform complexity was then summarised using a normalized isoform diversity (NID) metric. The NID score for *GBAP1* (0.0083) was higher than that of *GBA1* (0.0042), suggesting *GBA1* is under higher selective pressure. Moreover, entropy measurements confirmed that *GBAP1* exhibits higher entropy than *GBA1*, consistent with the greater isoform diversity observed above (**Supplementary Note 2**). Taken together, these results demonstrate that reference annotations systematically underestimate isoform diversity at this pseudogene-containing loci, and that population-scale indexing can recover biologically supported transcript structures that escape conventional annotation.

**Figure 4:**
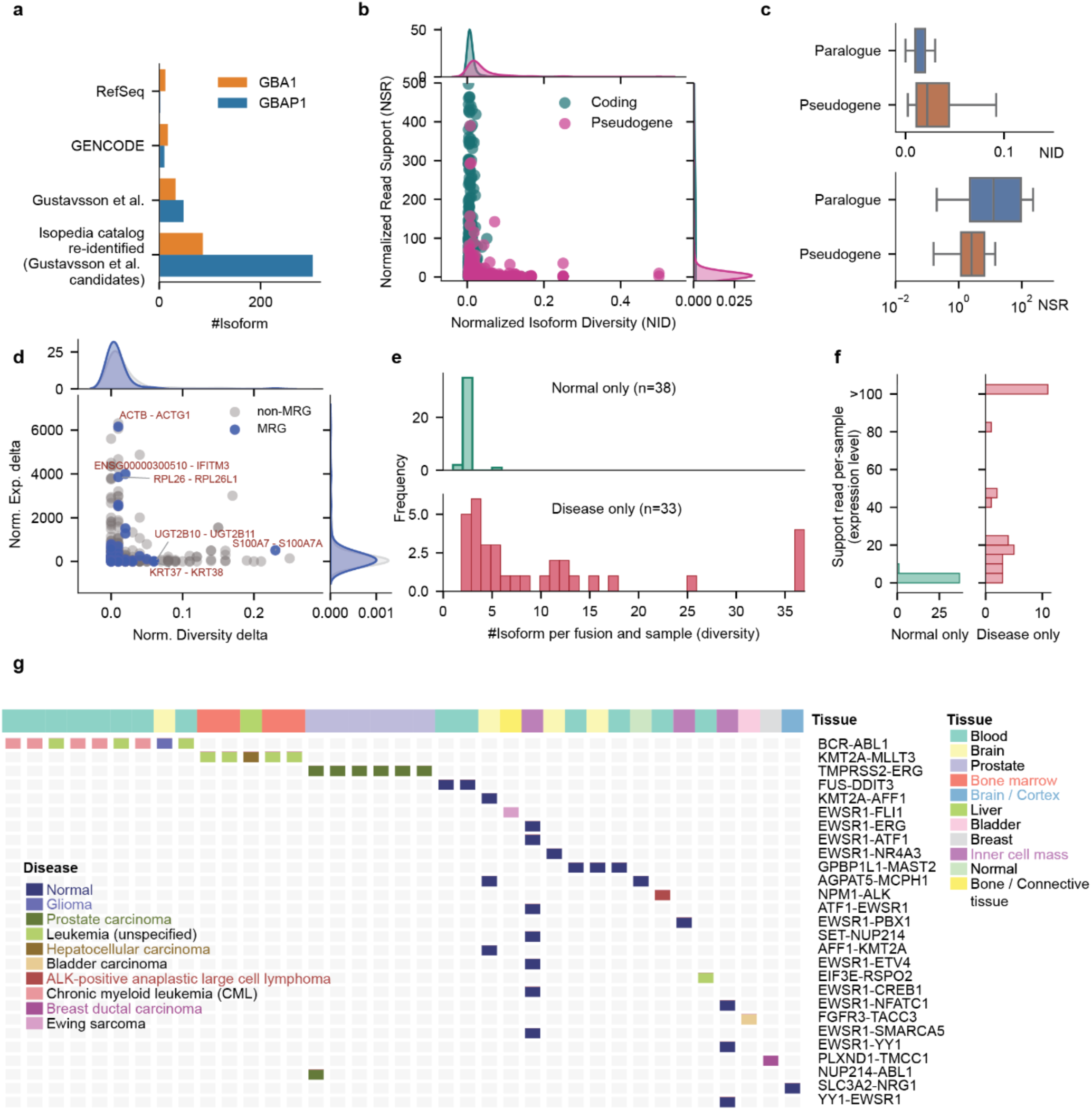
Isoform diversity revealed by the Isopedia catalog. **a)** Number of GBA1 and GBAP1 isoforms reported by different sources. The last category corresponds to isoforms re-identified in the Isopedia catalog when querying transcript candidates from Gustavsson et al. Twenty-nine samples overlapping with the original Gustavsson et al. study were excluded from the catalog, and only isoforms present in at least two of the remaining samples were retained. **b**) Normalized isoform diversity and expression of the pseudogene and its paired protein-coding gene. Top and right density plots summarize the distributions of NID and NSR (shown up to 500 for visualization), respectively. **c**) Comparison of NID and NSR between pseudogenes and paralogs. **d**) Normalized isoform diversity and expression of paralogous genes with 90-100% sequence identity. Top and right density plots summarize the distributions of NID delta and NSR delta, respectively. **e**) Isoform number comparison between normal-only and disease/cancer-only fusion events. **f**) Supporting read comparison between normal-only and disease-only fusion events. **g**) Fusion isoform matrix of normal-only and disease-only fusion events. **NID**: Normalized isoform diversity; **NSR**: Normalized support read.

To assess this genome wide, we next used Isopedia’s splice query to quantify the isoform diversity of pseudogene-coding gene pairs, assessing 1,471 pseudogenes with their matched 862 protein-coding genes. For this we used the same thresholds (at least two samples and two sequencing technologies) as before (see **Methods**). As we hypothesized, pseudogenes showed a higher NID (mean 0.04) compared to their corresponding protein-coding genes (mean NID 0.01) (**Fig. 4b**, **Supplementary Table S6**). Beyond isoform diversity, we also examined gene expression using a Normalized Support Read (NSR) score to assess whether pseudogenes differ from coding genes in both splicing complexity and transcriptional output. There was a clear separation of pseudogenes and protein-coding genes (**Fig. 4b**, **Supplementary Fig. S11**), highlighting not only higher isoform diversity among pseudogenes but also their lower expression (NSR: 9.70) in pseudogenes compared with NSR: 193.15 for paired coding genes (right density plot, **Fig. 4b**). This pattern of greater transcript diversification coupled with lower transcriptional output is consistent with relaxed selective constraint acting on pseudogenes.

Overall, the data support our hypothesis for pseudogene isoform diversity. However, it remained unclear whether this pattern arises from gene duplication itself or specifically from pseudogene formation. To disentangle these effects, we examined highly homologous paralog gene pairs (712 pairs, >90% identity), where the protein-coding copy is presumed functionally intact. Paralogues showed a markedly lower average NID of 0.02 compared to 0.04 for pseudogenes, and a substantially higher NSR of 259.87 versus 9.70 (**Fig. 4c**, **Supplementary Table S7**). These differences persisted even among more divergent paralogous copies (60–40% identity; NID 0.01, NSR 73.40; **Supplementary Table S18**). Thus, pseudogenes exhibit higher and likely less regulated isoform diversity coupled with lower expression, a pattern that cannot be explained by gene duplication alone.

Given the results on pseudogenes and paralogous genes, we investigated whether similar patterns could be observed in medically relevant paralogous genes. These are highly challenging to analyze as they often are highly similar or occur in complex regions^41^. We observed certain outliers, for example, *S100A7* vs. *S100A7A* showed a high delta on normalized splicing values (**Fig. 4d and Supplementary Table S7**). This may highlight their recent duplication and that one copy (*S100A7A*) is now under potentially reduced selection, which is important as both are reported to be medically relevant. In contrast, we observed *ACTB* vs. *ACTG1,* which showed a remarkable normalized isoform similarity but large expression. Collectively, these findings indicate that increased isoform diversity is not an inherent consequence of gene duplication but instead reflects a relaxation of regulatory constraint, suggesting that splicing complexity metrics may serve as a systems-level readout of functional divergence among paralogous genes. Indeed, understanding these dynamics carries translational significance, as it may help identify the dominantly expressed gene copy.

### Isoform-resolved fusion analysis reveals tissue-specific diversity and expression patterns

Gene fusions have been reported as important cancer drivers in multiple studies^42^. Consequently, we explored the use of Isopedia to prioritize gene fusions that are unique to a certain phenotype of the sample. More specifically, Isopedia can provide insights into the occurrence of a gene fusion and its prevalence in cancer and normal samples. To assess this, we leveraged the Isopedia catalog to annotate cancer-relevant gene fusions cataloged in COSMIC^43^ against the Isopedia catalog. Beyond breakpoint-level annotation, we characterized their full-length fusion isoform structures, an analysis that has rarely been performed as the functional diversity and regulatory complexity of fusion isoforms remain poorly characterized^44,45^. We queried the COSMIC fusion breakpoints against the Isopedia catalog. Among 1,314 fusion events queried, 112 were detected in the catalog, and we stratified them by disease annotation and tissue type. Of the detected fusion events, 41 (36.6%) (**Supplementary Table S9**) were observed in both normal and disease tissues from the Isopedia catalog, 38 (33.93%) were detected exclusively in normal tissues, and 33 (29.46%) were present only in disease tissues. The Isopedia fusion query revealed that fusions from disease tissues exhibited higher isoform diversity (**Fig. 4e)** compared to those from normal-labeled samples (10.82 vs 2.07 average isoforms per sample). For instance, the *BCR-ABL1* fusion had on average of 32.46 distinct isoforms across 9 cancer samples. Moreover, fusions exclusive to disease-labeled samples in the catalog showed substantially higher average read support (average of 148.91 reads/sample) compared to those exclusive to normal-labeled samples (∼2.07 reads/sample) (**Fig. 4f**). This was consistent with previous reports indicating that fusion transcripts associated with cancer can be detected in healthy tissues, typically at low abundance^46–48^. Overall, by leveraging Isopedia’s fusion query functionality, we demonstrated that COSMIC fusions, previously characterized only at the breakpoint level, harbor substantial isoform diversity. Fusions from disease-labeled versus normal-labeled samples in the catalog further exhibited markedly distinct profiles in both isoform complexity and expression abundance.

Beyond these global patterns, Isopedia re-identified well-established fusion-disease associations (**Fig. 4g**, **Supplementary Table S10**). The most frequently observed event (9 samples) was the *BCR-ABL1* fusion, a known hallmark and driver of chronic myeloid leukemia (CML)^49,50^. As expected, *BCR-ABL1* was predominantly detected in CML samples, with a single occurrence in a glioma sample. Isopedia also detected the *KMT2A-MLLT3* fusion, reported to be a driver event in leukemia^51^, in five samples, of which four were leukemia cases and one was a liver cancer case. In addition, the *TMPRSS2-ERG* fusion, commonly associated with prostate cancer^52,53^, was exclusively found in six prostate samples. Collectively, these analyses highlight that Isopedia characterizes gene fusions with respect to diseases and gives novel insights into their isoform diversity. This approach may facilitate systematic annotation of gene fusion events observed in normal tissues, helping to identify molecular precursors of cancer development.

## Discussion

In this study, we present Isopedia, a population-scale framework for indexing full-length transcriptomes from long-read RNA-seq data and enabling recurrence-based interpretation of RNA isoforms across tissues and studies. By aggregating 1,007 publicly available long-read transcriptomes into a unified and searchable catalog, Isopedia transforms isolated RNA-seq experiments into a cumulative resource for transcriptome interpretation. Importantly, this enables isoform discovery to be evaluated within an empirical population context rather than relative to a single, evolving reference annotation.

The maturation of isoform analysis now demands a fundamental shift in how RNA-seq data are interpreted. Despite major technical advances, interpretation remains constrained by reference-anchored paradigms that define isoform novelty relative to incomplete and inconsistently curated annotations. As a result, the field has remained in a discovery cycle, in which individual studies repeatedly report large fractions (26-70%) of “novel” isoforms^11,20,21^. By introducing transcript recurrence as a central organizing principle, Isopedia enables a transition from reference-dependent discovery to empirically grounded, population-aware interpretation, positioning transcriptomics alongside population genomics as a cumulative framework for biological inference. In our analyses, apparent novelty rates decreased by up to 26-fold (>95% annotated isoforms independent of sequencing technology) when evaluated against population-scale long-read evidence rather than reference annotations alone. Isoform frequency thus serves as an organizing axis to distinguish common transcript structures from rare or context-specific events, analogous to allele frequency in variant interpretation^54^.

Beyond annotation, population-scale transcript indexing reframes isoforms as quantitative population features rather than binary entities classified as “annotated” or “novel”. In this continuum model, transcript structures are interpreted according to their frequency and distribution across biological contexts, enabling refinement of reference catalogs and prioritization of rare or context-specific isoforms with potential functional relevance. Annotation thus becomes an evidence-weighted and evolving process rather than a static classification.

This population-aware framework enables biological insights that are not accessible in single-study analyses. We find that pseudogenes exhibit elevated isoform diversity coupled with reduced expression, consistent with relaxed regulatory constraint, whereas paralogous protein-coding genes maintain constrained transcript architectures despite high sequence similarity. These observations suggest that isoform diversity can act as a systems-level proxy for regulatory constraint. Similarly, fusion transcript analyses reveal that canonical gene fusions comprise structurally heterogeneous isoform ensembles, with disease-associated fusions showing increased diversity and expression relative to those observed in normal tissues. Together, these findings demonstrate how population-aware transcriptomics enables systematic investigation of transcript regulation, evolution, and disease relevance. Importantly, this framework also has direct translational implications. By establishing a population baseline for transcript structure and abundance, population-scale indexing enables prioritization of rare splicing events and fusion transcripts that distinguish physiological from pathological states. This provides a foundation for improved functional variant interpretation, biomarker discovery, and therapeutic target identification, particularly in diseases where transcript-level variation contributes to mechanism.

A current limitation is that the catalog reflects the uneven distribution of publicly available long-read datasets across tissues and disease states. However, this is a constraint of data availability rather than the framework itself; as sequencing expands across diverse populations and clinical contexts, the resolution of this population-native index will increase. Importantly, the current inclusion of substantial cancer transcriptomes enables immediate identification of cancer-associated isoforms and fusions. Researchers can now evaluate whether candidates are recurrent across malignancies, tissue-restricted, or present in healthy samples.

Overall, Isopedia establishes a population-native foundation for transcriptome interpretation. By transforming isoform discovery from an episodic, annotation-dependent exercise into a cumulative and context-aware process, it enables transcript diversity to be interpreted with a rigor comparable to that achieved in population genomics. As datasets from underrepresented tissues, ancestries, and developmental stages grow, the interpretive power of our population-aware indexing will continue to improve, further reducing the search space for functionally relevant transcript variation. Thus, isopedia will be essential for converting transcript variation into robust biological and clinical insight.

## Methods

### Isopedia workflow

Isopedia is designed to index large-scale datasets and provide efficient retrieval and query functionality. Its commands are organized into two categories. For indexing, Isopedia provides three subcommands: profile, which extracts isoform structural signatures from individual long-read RNA samples; merge, which consolidates isoform signatures across multiple samples into a unified archive file; and index, which constructs a B-plus tree index for efficient lookup. For querying, Isopedia provides three subcommands corresponding to different input types: isoform structure queries, individual splice junction queries, and fusion queries. The pan-transcriptome isoform representation (PTIR) is the core data structure that enables Isopedia to construct compact catalogs and perform rapid query comparisons.

#### Population-native isoform representation

Isopedia is built upon a core concept and data structure termed pan-transcriptome isoform representation (PTIR). A PTIR represents an aggregation of long-read transcriptome alignments that share an identical splice junction chain. For each read contributing to a PTIR, read-specific mapping information is retained to enable reconstruction of the full alignment status. This information includes the start, end, and strand of the primary alignment, as well as the genomic coordinates and strand of any supplementary alignment segments.

PTIRs are designed to support aggregation of reads originating from different samples, provided that they share the same splice junction structure. For each PTIR, two sample-indexed vectors are maintained to store the pointer and length of sample-specific elements within the aggregated record. This design allows PTIRs to be constructed from a single sample and incrementally merged across multiple samples with minimal overhead, forming the basis for population-scale isoform representation.

#### Profiling isoforms in individual samples

Isopedia profiles isoform signals by scanning raw alignment records from individual samples. By default, only primary alignments are considered for splice junction extraction. Splice junctions, along with the genomic start and end positions of each read, are parsed from the CIGAR string (e.g. refskip N is used to determine the intron boundary) of the primary alignment. For supplementary alignments, only the genomic start and end positions of each aligned segment and the corresponding strand information are retained. Reference-span positions derived from supplementary alignments are additionally collected to support downstream fusion gene detection.

Initially, each read is assigned to a PTIR based on its primary splice junction chain. Specifically, the ordered vector of splice site positions is hashed (ahash library^55^) to produce a unique signature for each distinct isoform structure. Reads that share an identical hash signature, and thus the same splice junction configuration, are merged into a single PTIR. This approach enables efficient comparison and consolidation of reads at the single-sample level.

#### Merging isoform profiles across multiple samples

To construct population-scale PTIRs, Isopedia takes as input the compressed single-sample profiles from multiple samples. A manifest file is required to specify sample identifiers, paths to single-sample profile files, and associated metadata. PTIRs from different samples are merged by comparing their splice junction hash signatures, allowing isoforms with identical structures to be aggregated across samples.

To limit memory usage during this process, Isopedia employs a min-heap-based strategy to iteratively merge PTIRs from multiple samples. The final output is a single compressed archive file containing PTIRs spanning all samples. This file is ordered by chromosome and the genomic position of the first splice junction within each PTIR to reduce I/O overhead during downstream querying. In addition, a temporary position-pointer index is generated, recording the genomic position of each splice site and the corresponding pointer of the PTIR record within the aggregated file. This index serves as the input for the subsequent indexing step.

#### B-plus tree indexing

Population-scale indexing generates a large number of PTIRs, necessitating efficient retrieval of specific transcripts. To address this, we implemented a B-plus tree index that maps the genomic coordinates of each isoform to the corresponding PTIR address (PTIR pointer) within the archive file. Because all position-pointer pairs are pre-sorted and no new records are introduced during indexing, the tree can be constructed bottom-up starting from the leaf nodes, ensuring a fully populated tree structure.

A separate B-plus tree file is generated for each chromosome. The indexing process supports multi-threaded execution to accelerate construction. The manifest file is required at this stage to associate PTIR records with sample-level metadata, enabling efficient retrieval of population-level isoform information during query operations.

#### Querying the index

Isopedia employs a unified query schema for accessing the population-scale catalog. All queries are first decomposed into a set of genomic positions, optionally extended by a user-defined number of flanking base pairs (specified by the -f option; default 10 bp). This flanking tolerance accommodates alignment uncertainty at splice boundaries commonly observed in long-read RNA sequencing, analogous to the splice-site correction windows employed by long-read transcript assembly tools such as EXPRESSO^56^. Moreover, as a query tool, Isopedia operates on user-supplied transcript models (GTF file). For each query, the corresponding chromosome-specific B-plus tree is loaded, and a set of candidate pointers pointing to PTIR records is retrieved from the tree.

The returned PTIR pointers are subsequently post-processed according to the query type to identify valid target records. For isoform queries, which consist of a list of genomic positions that derived from GTF file, a PTIR is considered a valid match only if its pointer matches up all splice junction configuration. For splice junction queries, which are defined by two genomic positions corresponding to the start and end of a junction (ie. BED-like files), any PTIR pointer matches both coordinates is considered a valid match. For fusion queries, which are two coordinates (BED-like file), all returned PTIR pointers are treated as candidate records and are further examined by inspecting their supplementary alignment information to determine whether they overlap the queried genomic regions. Notably, all three query functions allow wobble match coordinates which are enabled by -f option.

Once the final set of PTIR pointers has been determined, Isopedia loads the corresponding PTIR records from disk to retrieve query-type-specific information, such as sample support, read counts, and alignment structures. The output of all three query functions is a table with each row representing a query entry, and columns representing the supporting read count or expression value in TPM for each sample in the Isopedia catalog.

The compact PTIR representation enables Isopedia to operate within constrained memory environments (for example, 16 GB RAM for isoform queries). To enable efficient streaming access, Isopedia requires that input queries be pre-sorted by chromosome name and genomic position. Each chromosome-specific B-plus tree embeds a least-recently-used (LRU) cache to retain recently accessed position-pointer mappings, thereby reducing disk I/O and accelerating repeated or spatially localized queries.

Quantification of queried transcript with expectation-maximization algorithm

Reads sharing identical splice junctions are aggregated into a single representative unit, thereby reducing storage requirements and computational overhead. This aggregation naturally defines equivalence classes^57,58^ for efficient transcript quantification. We adopt the expectation-maximization (EM) algorithm to estimate transcript-level abundance from these equivalence classes. A merged splice junction class corresponds to one PTIR and contains the set of all transcripts compatible with that aggregated read. For each independent sample *s*, the compatible transcript set of PTIR *m* is denoted *s_m_*, and its read support in sample *s* is *c_m,s_*, the likelihood function of the abundance can be modeled as:

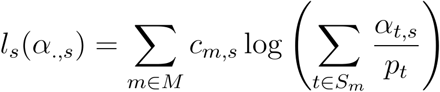

We maximize *l*_*s* for each sample independently, since PTIR memberships are shared but abundances are sample-specific. To correct for transcript-specific sampling opportunities, we compute *p_t_* based solely on transcript splice junction complexity. For each transcript *t*, *J_t_* denotes the number of splice junctions, and *p_t_* = *J_t_* + 1. We define *p_t_* = *J_t_* + 1 to avoid a value of zero for single-exon transcripts. We use the EM algorithm to estimate the expected read counts of transcripts. Briefly, we define gamma as the latent variable to indicating the probability that a read belongs to transcript *t* in the E-step:

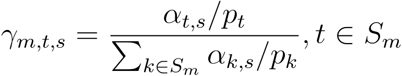

In the M-step:

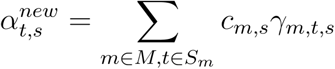

In practice, PTIR are shared across samples, and EM updates are performed independently for each sample (implemented in a vectorized manner across samples). If a PTIR has zero supporting reads in sample *s* (i.e., *c_m,s_* = 0), the abundance estimation ignores the PTIR. Transcripts per million (TPM) is used to represent transcript abundance. In the long-read RNA sequencing context, TPM is identical to counts per million (CPM), as long reads capture full-length transcripts and thus do not require normalization by transcript length^15^.

### Building a catalog of 1,007 human transcriptomes

The datasets were downloaded either in BAM format or as raw reads. All reads were aligned to the GRCh38.p14 reference using Minimap2^59^ (2.20-r1061) with the parameter “-ax splice:hq -uf” for PacBio datasets and “-ax splice -uf -k 14” for ONT datasets. Raw signals were extracted from the BAM files using the Isopedia profile command with default parameters. Isopedia merge and Isopedia index were used to aggregate independent files and create indexes, respectively, using default parameters. The metadata for each dataset were collected along with the datasets and manually curated and standardized. The curated meta data table can be found at **Supplementary Table S1**.

### Benchmarking

#### Simulated datasets

The simulated benchmark, which consists of two datasets for PacBio and Oxford Nanopore technologies, was directly downloaded from the LRGASP^19^ website. Reads were simulated according to a subset of transcripts from the GENCODE V38 annotation by LRGASP. In total, 48,192 and 47,902 transcripts were simulated for PacBio and ONT, respectively.

#### Re-identification of isoforms using simulated datasets

Simulated datasets were obtained from the LRGASP project^19^. The paired reference annotation (based on GENCODE V39) and reference genome (based on GRCh38) from LRGASP were also used for benchmarking. The ground truth consists of a list of transcript IDs (from the LRGASP annotation) for PacBio and ONT, respectively. Minimap2 was used to align simulated reads against the LRGASP reference genome with platform-dependent parameters (PacBio: -ax splice:hq -uf; ONT: -ax splice -uf -k14). All tools, including Isopedia, were run with their default parameters and provided with the LRGASP reference annotation and reference genome as input where required. The transcripts reported by each tool were compared to LRGASP reference annotation using GFFCompare^37^ (v0.12.9) and then evaluated against ground truth. Based on this, we define: True Positive (TP) = transcripts in the ground truth that were detected; False Positive (FP) = detected transcripts not in the ground truth; False Negative (FN) = transcripts in the ground truth that were not reported. Recall is defined as TP/(TP+FN), Precision = TP/(TP+FP), and F1 score = 2 × (Precision × Recall) / (Precision + Recall).

##### Stringtie2

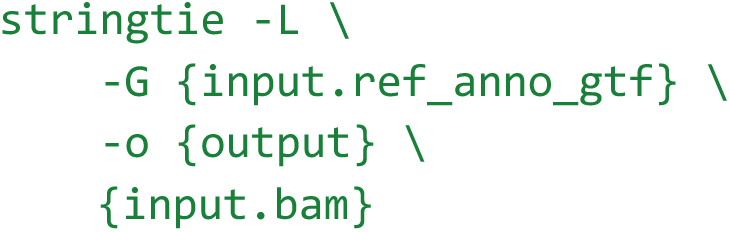

##### Bambu

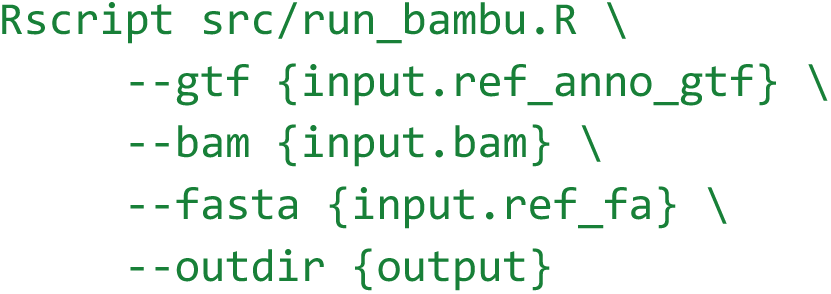

##### Isoquant

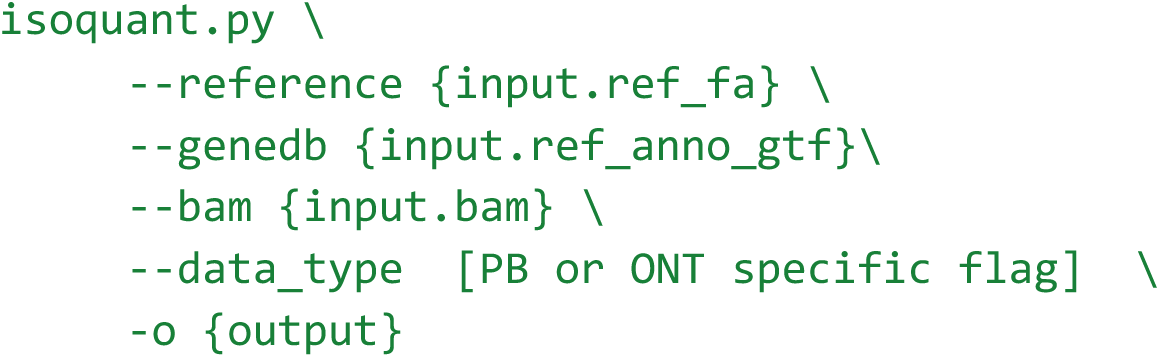

##### Oarfish

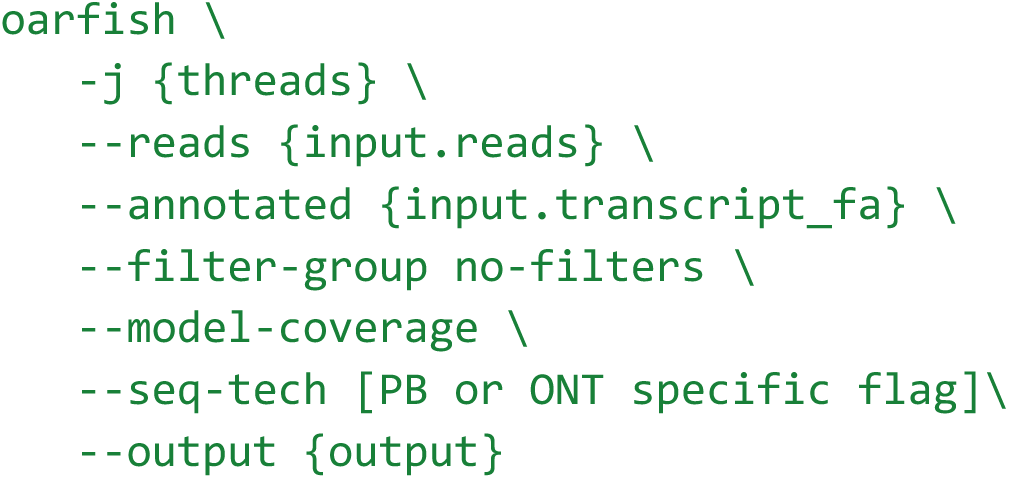

#### Quantification comparison using simulated datasets

Ground-truth TPM values were obtained from the file hs_GENCODE38.basic_annotation.counts.tsv.gz provided by the LRGASP. TPM estimates were extracted from the output of each tool. Comparisons were restricted to simulated isoforms from the PacBio and ONT datasets, respectively. Spearman correlation was used to assess consistency between the ground truth and individual tools, as well as between pairs of tools.

#### Consistency evaluation using real sequencing datasets

We evaluated Isopedia using 18 real sequencing datasets from the LRGASP WTC11 cell line, representing six combinations of library preparation methods (CapTrap, cDNA, R2C2, dRNA) and sequencing platforms (PacBio, ONT), with three biological replicates each. SIRV-Set4 transcripts were spiked into all samples as controls. Isopedia indexed all datasets using the GENCODE v38 annotation (237,012 transcripts) as the reference. Re-identification robustness was assessed by the consistency (proportion of transcripts detected in all or none of the replicates) rate across replicates. Briefly, each tool was run on all 18 datasets individually, and the results were grouped by replicate. For each transcript across the three replicates of a given library preparation method, detection in either all three or none of the replicates was considered consistent; otherwise, it was counted as inconsistent. Similarly, quantification stability was measured using the coefficient of variation (CV) of mean expression across all transcripts among replicates. Performance on spike-in controls was evaluated by calculating recall rates for SIRV transcripts.

### HG002 transcriptome analysis

BAM files for PacBio, ONT, and Illumina sequencing (mRNA) were downloaded directly from the GIAB website. Each technology includes three BAM files (“gm24385”, “gm26105”, “gm27730”). Because the PacBio BAM files were unaligned, we used minimap2 to align them to the GRCh38 reference with parameters -ax splice:hq -uf. Illumina reads were aligned to the reference genome using STAR^60^ (v2.7.9a) with default parameters. We used StringTie^16^ (3.0.2) to identify transcripts for all three technologies, guided by GENCODE V49. Particularly, for PacBio and ONT BAM files, the -L parameter was used. StringTie merge was used to merge the three GTF files within each technology to generate a unified GTF per technology with default parameters. Transcripts lacking strand information were excluded, as GFFCompare requires it for precise comparison. Intersections of transcripts across technologies were compared using GFFCompare with default parameters; class code ‘=’ was considered indicative of identical transcripts. Transcripts from each technology were annotated against GENCODE and RefSeq using GFFCompare. Isopedia annotation was conducted using the following command: isopedia isoform -g {query.gtf} -o {out.isoform.gz} -i {isopedia_catalog}.

### Querying GENCODE and RefSeq annotations

The GENCODE release V49 basic annotation GTF file was obtained from the GENCODE official release website. The RefSeq release 110 annotation GTF file was obtained from the NCBI official website. Both GTF files were used as input for the Isopedia isoform query with the following command:

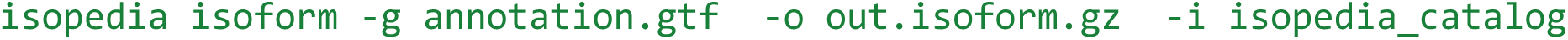

### Pseudogene and paralog analysis

To assess the isoform diversity of GBA1 and GBAP1 at population scale, we collected their reported isoform numbers from different sources: *GBA1* and *GBAP1* transcript counts from GENCODE and RefSeq were obtained from their respective GTF files. The counts reported by Gustavsson et al^40^. were taken from the main text of the paper. To re-identify transcript candidates of *GBA1* and *GBAP1* that reported by Gustavsson et al., we queried transcript candidates (*GBA1*:2,368, *GBAP1*:3,083) from Gustavsson et al. against the Isopedia catalog to assess how many of them could be independently re-identified. We removed the candidates on the plus strand as both *GBA1* and *GBAP1* are on the minus strand. The mono-exon transcripts were also excluded. Briefly, transcript candidates in GTF format from the same study were queried against the Isopedia catalog using the following command:

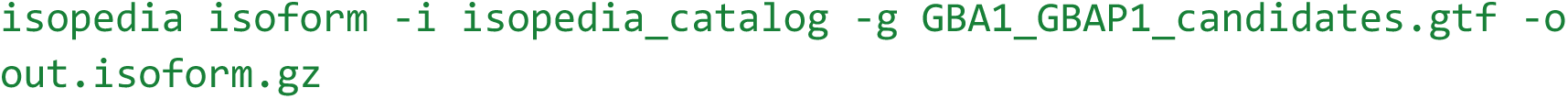

Because the Isopedia catalog includes 29 samples from Gustavsson et al.^40^, we excluded these overlapping samples from the isopedia catalog to avoid validation biases. We further required that queried transcript candidates be present in at least two of the remaining samples to ensure confident validation.

For the genome wide scan, we build up a catalog to query all pseudogenes. We applied a sequence-similarity-based approach to build the pseudogenes list. Briefly, genes and their associated transcripts annotated as “transcribed_*_pseudogene” in the GENCODE V49 were extracted as pseudogene candidates. Protein-coding genes and their transcripts were also extracted as coding-gene candidates. Transcript sequences for both pseudogene and coding-gene candidates were retrieved from the corresponding transcript sequences in GENCODE V49. Minimap2 was then used to align pseudogene candidate sequences to protein-coding gene sequences with the parameter set “-ax asm20”. A pseudogene candidate was retained if it was primarily mapped to the corresponding protein-coding gene on either the positive or negative strand. The splice junctions of all eligible candidate transcripts were converted into BED format and processed using the Isopedia splice mode. A returned record was retained if it was supported by at least two samples and both PacBio and ONT technologies. Total supporting reads and isoform count obtained from Isopedia per gene and normalized are used to report expression level and isoform diversity. Briefly, the normalized support reads (NSR) for gene i is defined as:

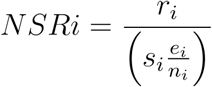

The normalized isoform diversity (NID) is defined as:

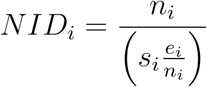

Where *r* denotes the total supporting reads, *e* denotes the total exon count from all isoforms detected, *s* is the number of positive samples, and n is the number of isoforms.

For paralogs analysis, we downloaded the paralogue gene from ENSEMBL biomart with the GRCh38.p14 reference version. Only entries annotated as ‘within_species_paralog’ were selected for downstream analysis. We restricted paralogue genes to those with a percentage of identity between 90% and 100% and 40-60%, respectively. The splice junctions of all eligible candidate transcripts were converted into BED-like format and processed using the Isopedia splice mode with command:

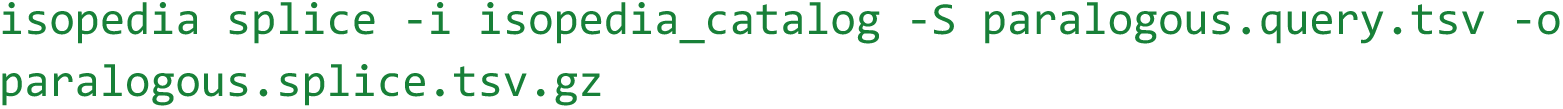

A returned record was retained if it was supported by at least two samples and both PacBio and ONT technologies. The NID and NSR calculated as described above.

### COSMIC fusion analysis

Cosmic fusion was downloaded from COSMIC^43^ with version 103. The raw fusion records were processed by customized scripts to extract the reported fusion breakpoints. A BED-like file was generated for Isopedia. Isopedia queried fusion breakpoints file with the following command:

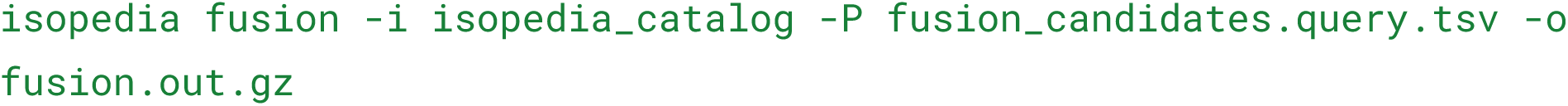

For each detected fusion event, isoform diversity was defined as the total number of distinct isoform structures observed across both gene partners, and average diversity was calculated by dividing this count by the number of positive samples. Similarly, fusion read support was determined by summing the supporting reads across all fusion isoforms, and average read support was obtained by dividing the total read support by the number of positive samples.

## Data availability

Isopedia catalog has been hosted at ftp://hgsc-sftp1.hgsc.bcm.tmc.edu//rt38520/isopedia_index_hs_v1.0.tar.xz. The manifest of all samples is available at **Supplementary Table S1.**

LRGASP datasets were downloaded from: https://www.gencodegenes.org/pages/LRGASP/. The HG002 BAM files were downloaded from https://ftp-trace.ncbi.nlm.nih.gov/ReferenceSamples/giab/data_RNAseq/AshkenazimTrio/HG002_NA2438 5_son. The subdirectories are “Baylor_PacBio” for PacBio, “Mason_ONT-directRNA” for ONT, and “Google_Illumina/mRNA” for Illumina.

GENCODE annotation was downloaded from https://www.gencodegenes.org/human/releases.html. NCBI RefSeq was download from https://ftp.ncbi.nlm.nih.gov/refseq/H_sapiens/annotation/GRCh38_latest/refseq_identifiers/ The paralogous gene table was downloaded from https://useast.ensembl.org/info/data/biomart/index.html. Fusion breakpoints were collected at https://cancer.sanger.ac.uk/cosmic/download/cosmic/v103/fusion. Medical relevant gene list was obtained from https://github.com/usnistgov/cmrg-benchmarkset-manuscript/blob/master/data/gene_coords/unsorted/GRCh38_mrg_full_gene.bed

## Code availability

The source code and binaries of Isopedia is available at https://github.com/zhengxinchang/isopedia.

## Acknowledgement

This research was supported in part by the National Institute of Health (1R01HG011774-01A1, 1U01HG011758-01).

## Supplementary Figures

**Supplementary Fig. S1:**
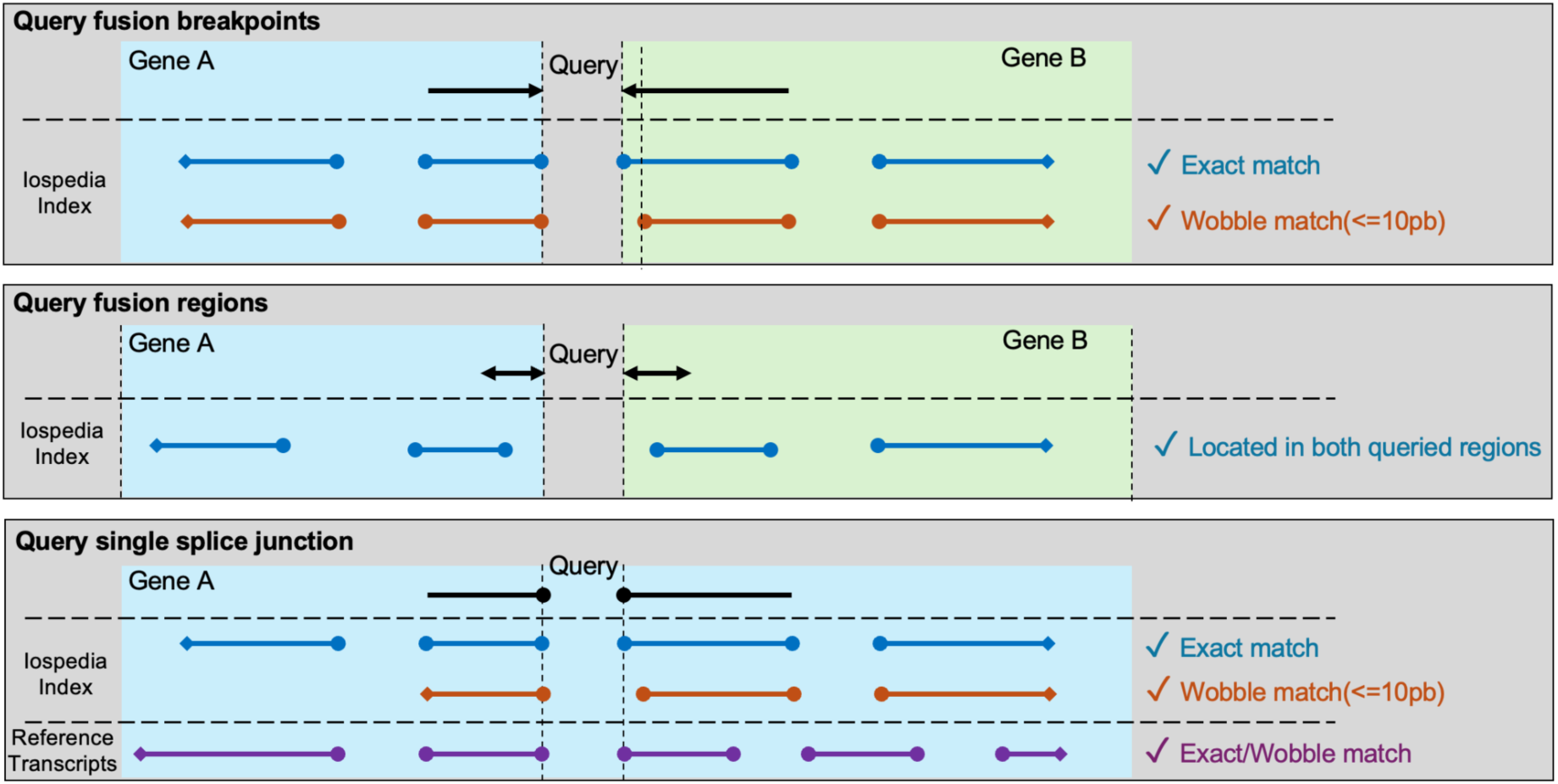
Overview of how Isopedia examines candidate reads. From top to bottom: query fusion breakpoints, query gene regions for fusion discovery, and query single splice junctions.

**Supplementary Figure S2:**
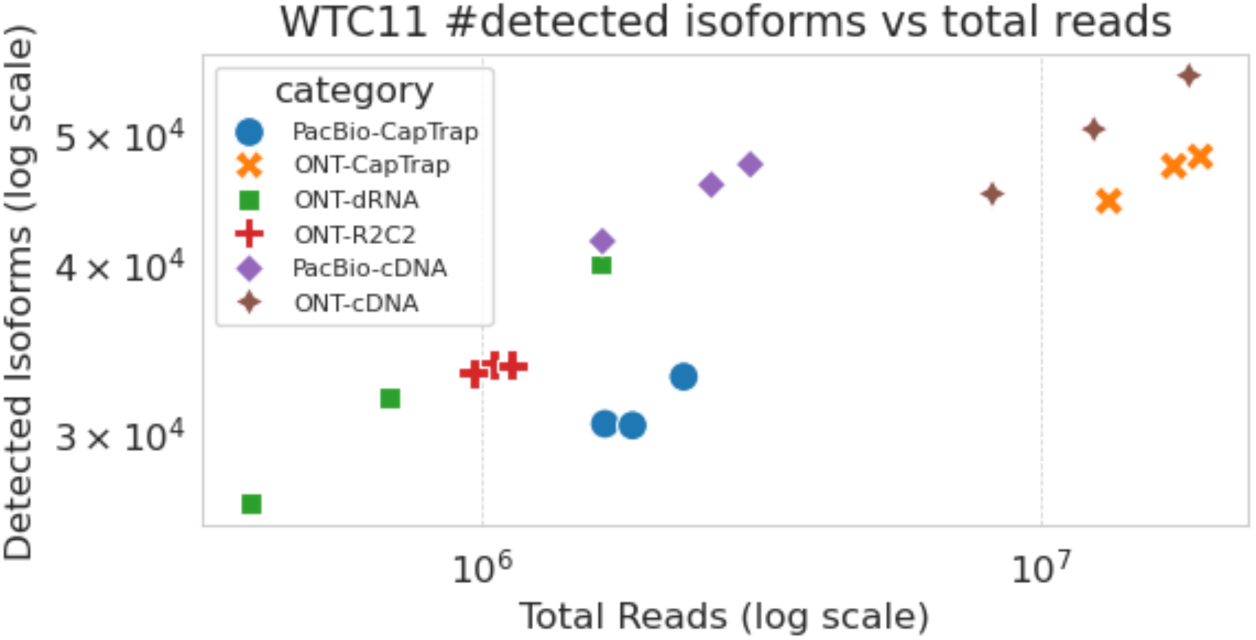
Detected isoform numbers versus total read counts in 18 WTC11 datasets across six categories. Each dot represents one dataset, and both color and shape indicate the corresponding library preparation method and sequencing technology.

**Supplementary Figure S3:**
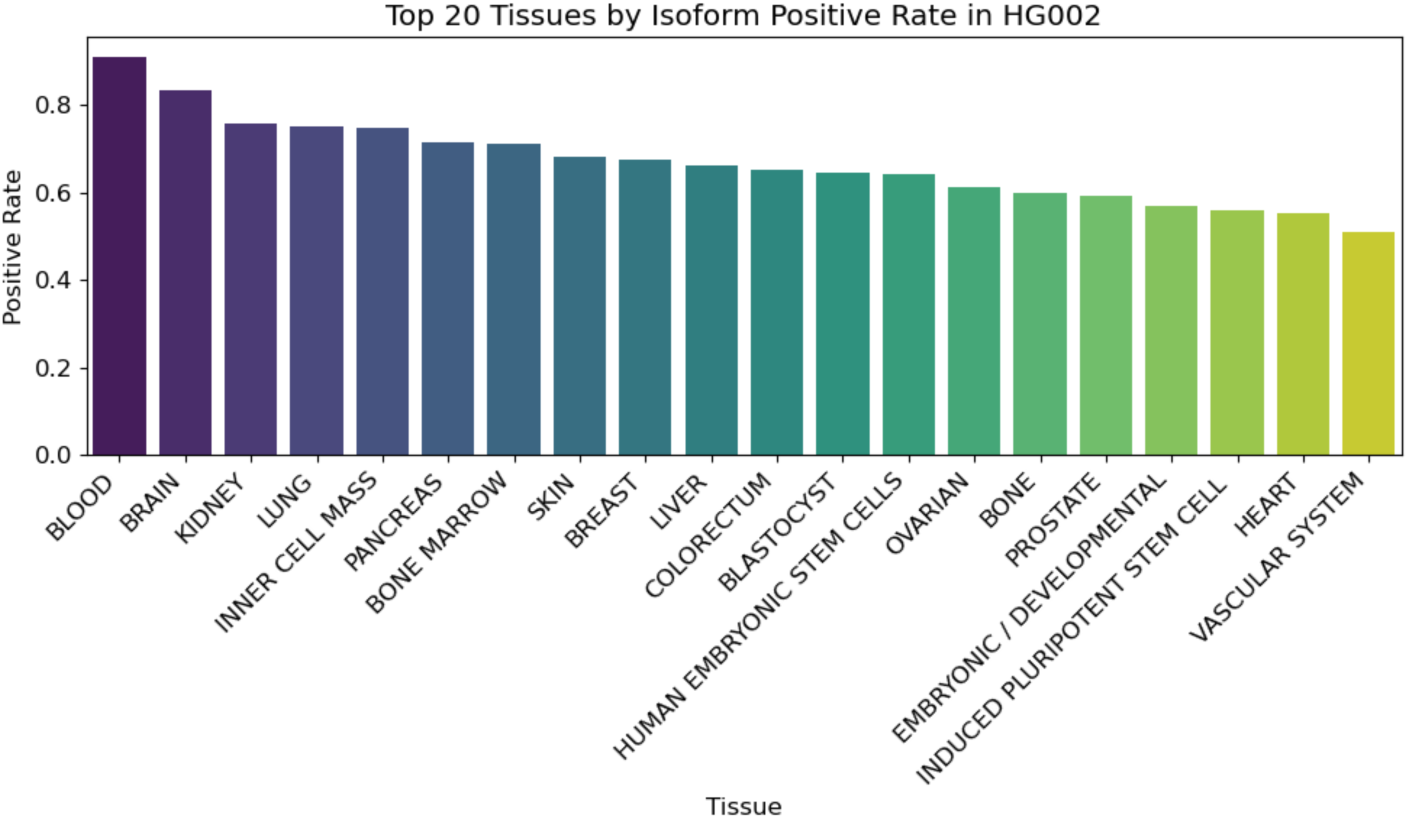
Percentage of HG002 isoforms were found in Isopedia catalog with different tissues.

**Supplementary Figure S4.**
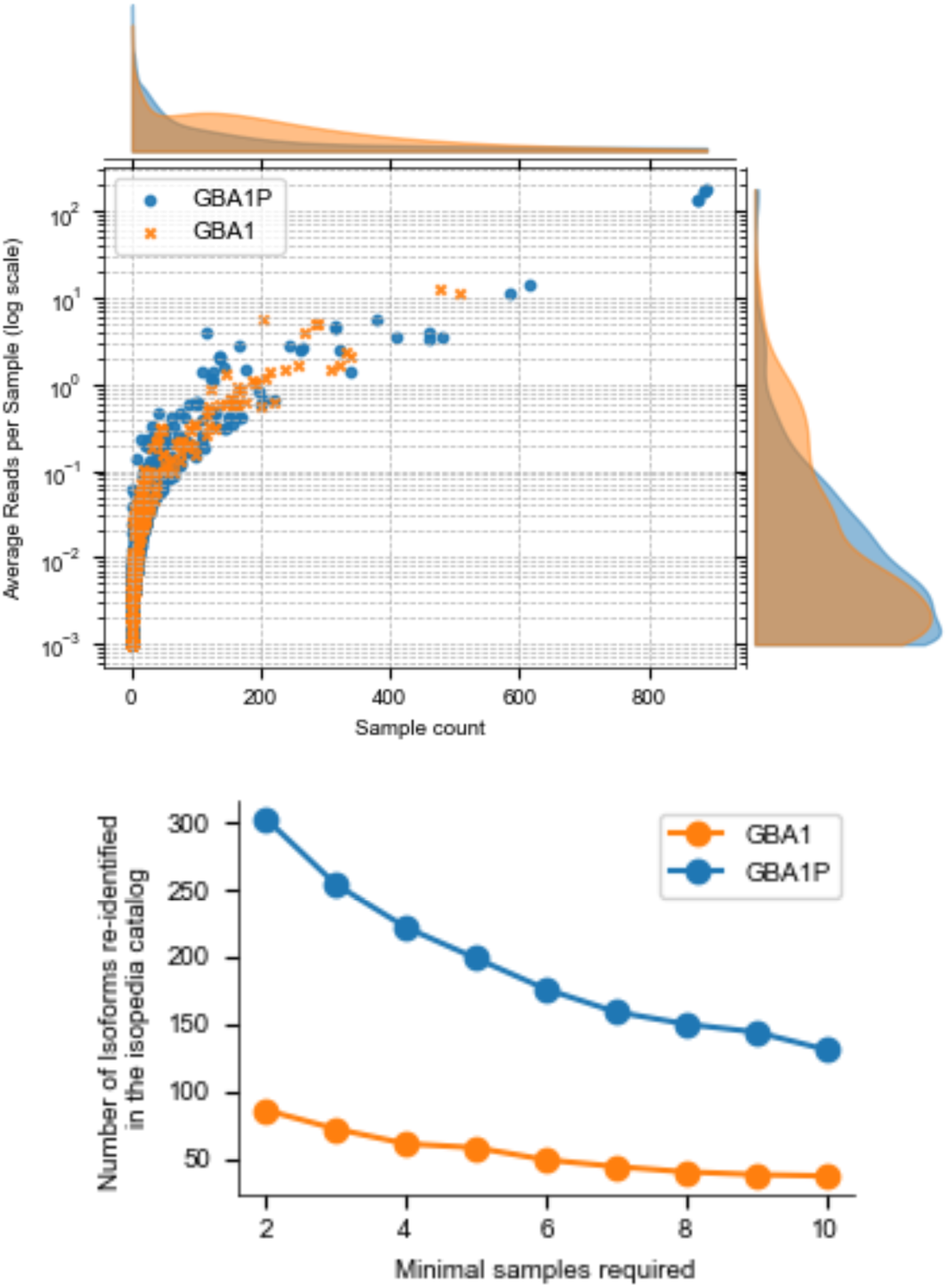
GBA1 and GBAP1 (pseudogene) transcripts identified in the Isopedia catalog. Upper subfigure, The X-axis represents the sample count in log scale, and the Y-axis shows the average read per sample among all positive samples. Isoforms are derived from Gustavsson et.al. transcript candidates. Lower figure, the GBA1 and GBAP1 transcripts (y-axis) re-identified by isopedia catalog with different minimal sample required cutoffs (x-axis).

**Supplementary Figure S5.**
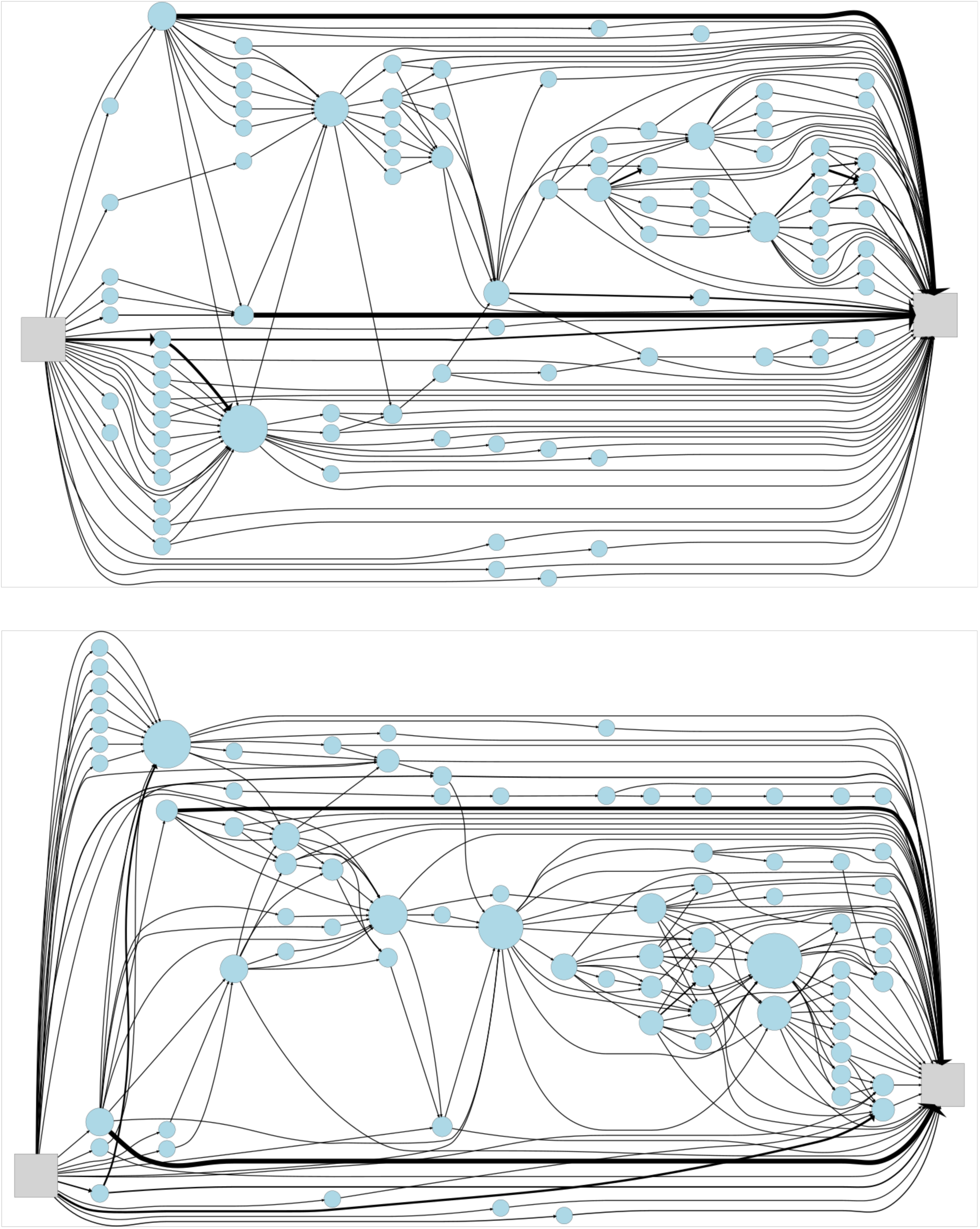
Isoform graph of GBA1 and GBAP1. The isoforms were obtained from Isopedia catalog by querying all splice junctions of all isoforms from GENCODE V49 of GBA1 and GBAP1.

**Supplementary Figure S6.**
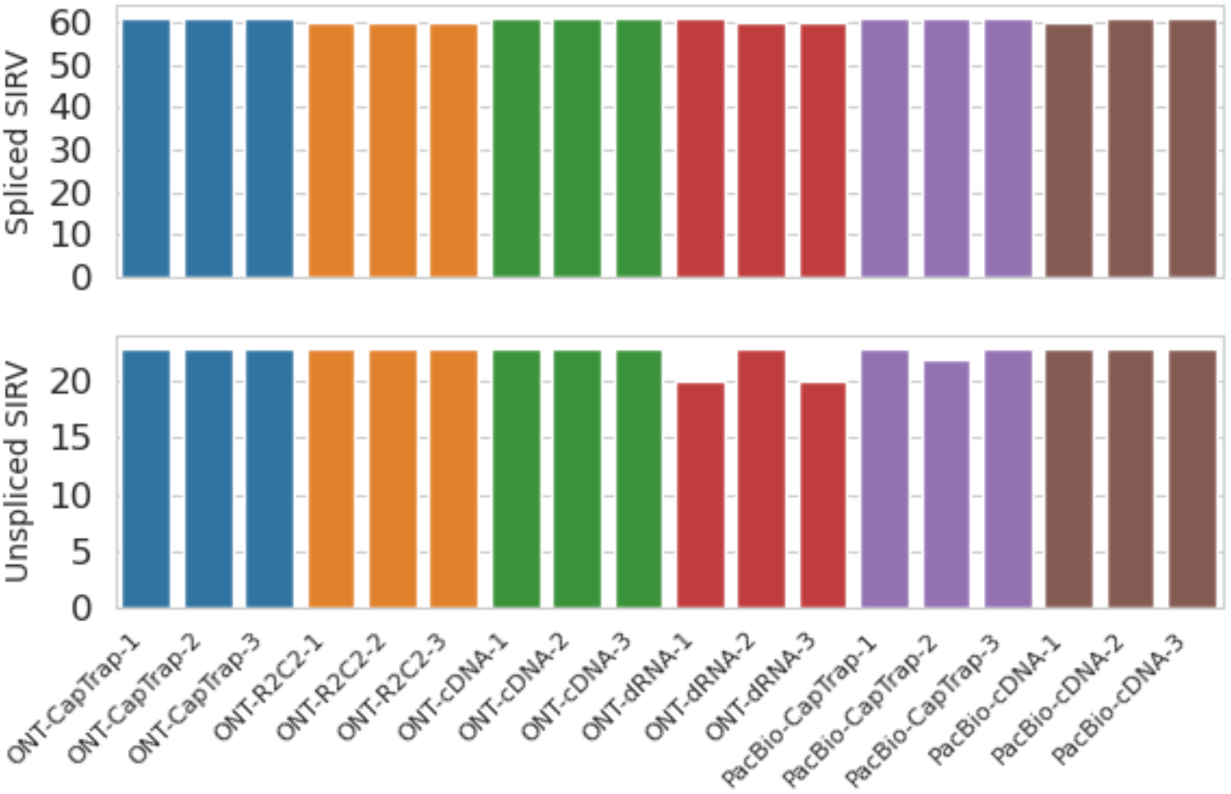
Recall of SIRV spike-in transcripts across all WTC11 datasets. The top panel shows transcripts containing at least one splice junction, whereas the bottom panel shows transcripts without splice junctions. We evaluated the re-identification of isoform performance of Isopedia on real datasets by assessing the recall of SIRV transcripts. For the 61 spliced SIRV transcripts, Isopedia identified all transcripts in 12 samples and 60 out of 61 transcripts in the remaining six samples. In particular, all 61 transcripts were consistently recovered across all replicates from ONT-CapTrap, PacBio-CapTrap, and ONT-cDNA libraries, whereas the other library types contained at least one sample missing a single transcript. For the 23 unspliced (mono-exonic) SIRV transcripts, Isopedia successfully identified all transcripts in 15 of the 18 samples. In the remaining cases, 20 out of 23 transcripts were recovered in two ONT-dRNA replicates, and 22 out of 23 transcripts were recovered in one PacBio-CapTrap replicate (**Supplementary Table S11**).

**Supplementary Figure S7.**
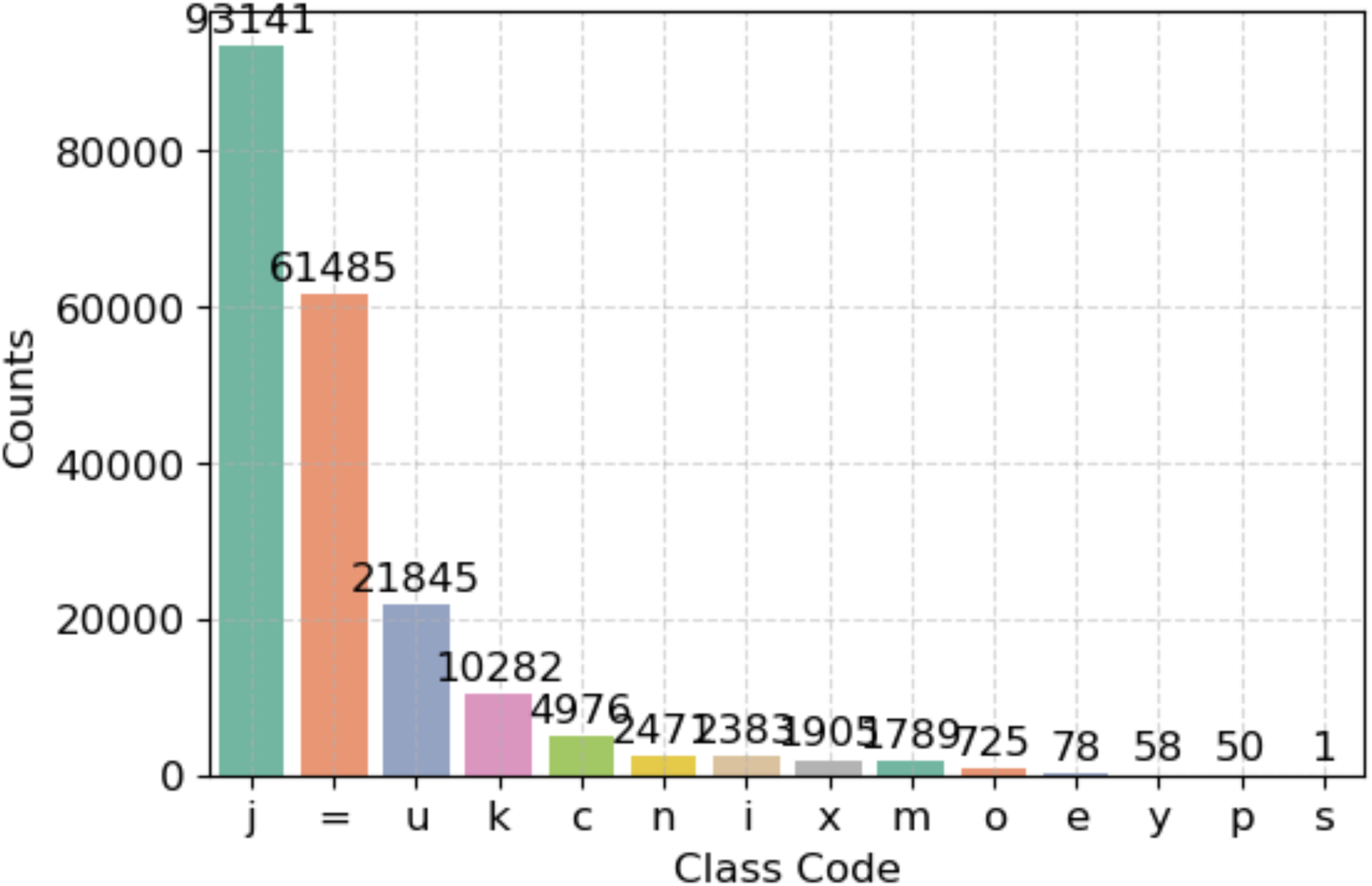
Transcripts comparison between GENCODE and NCBI RefSeq

**Supplementary Figure S8.**
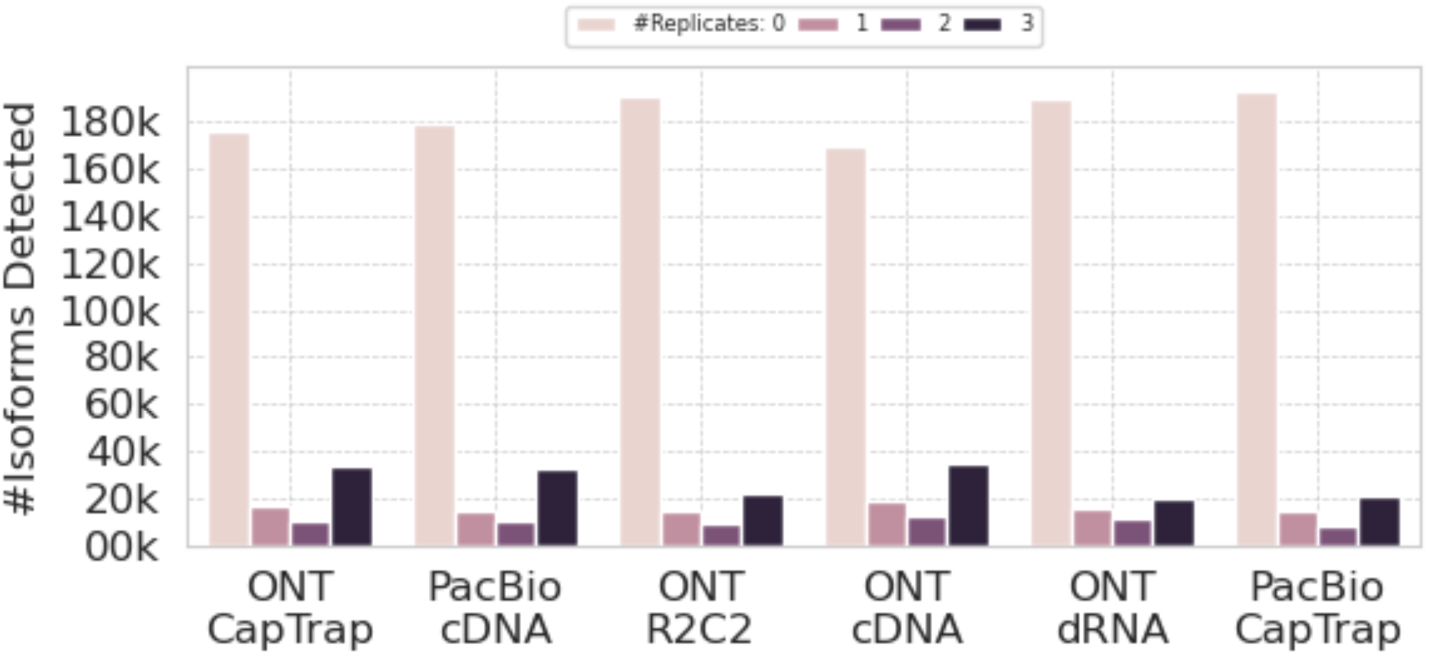
Consistency assessment of transcripts reported by Isopedia across replicates for each library preparation and sequencing technology combination. Isoforms that are either absent from all three replicates or present in all three replicates are considered consistent; isoforms detected in only a subset of replicates are classified as inconsistent.

**Supplementary Figure S9.**
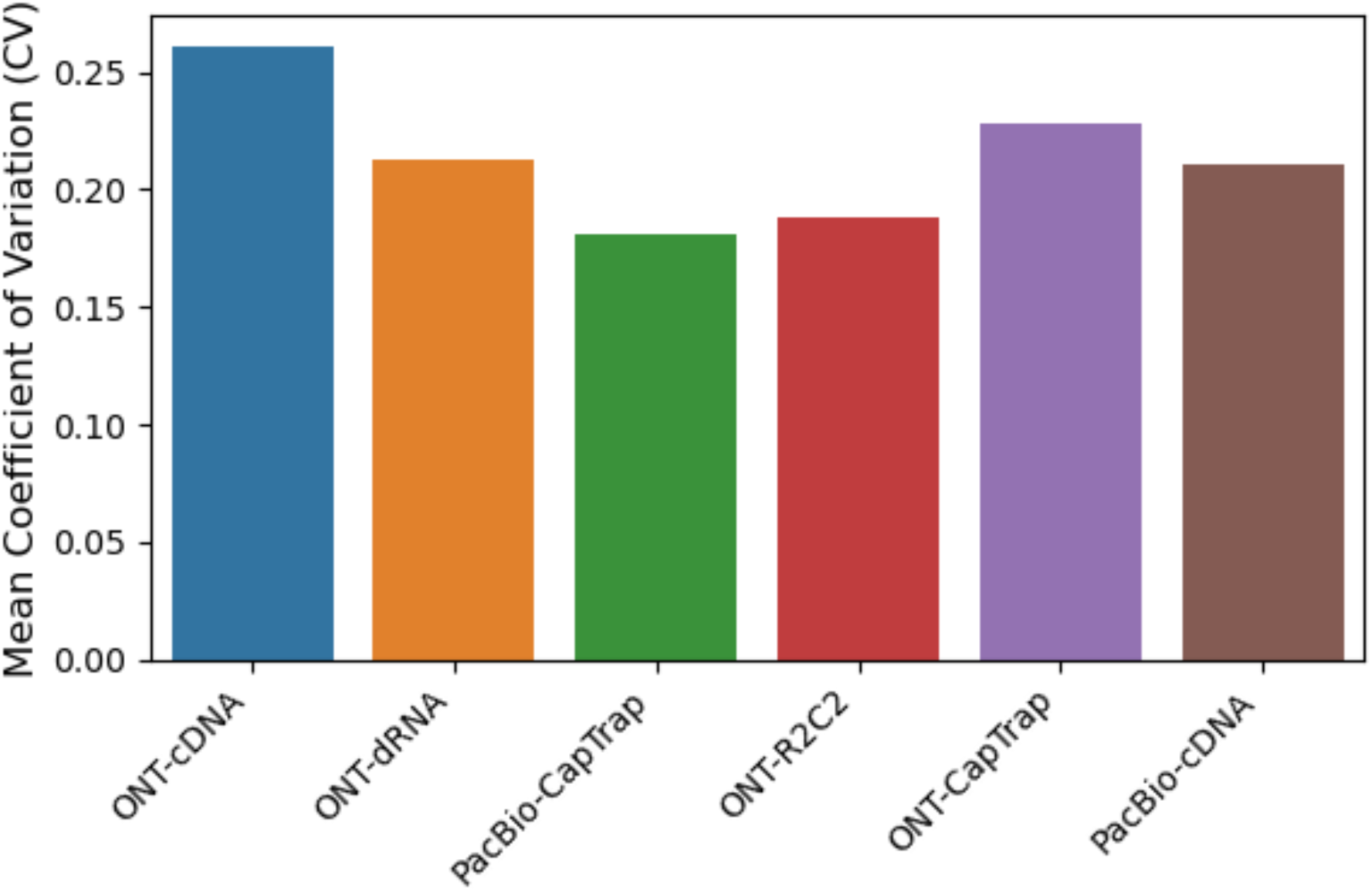
Mean coefficient of variation of transcripts from the Isopedia queried by GENCODE V49 annotation across three replicates for each library preparation and sequencing technology combination.

**Supplementary Figure S10.**
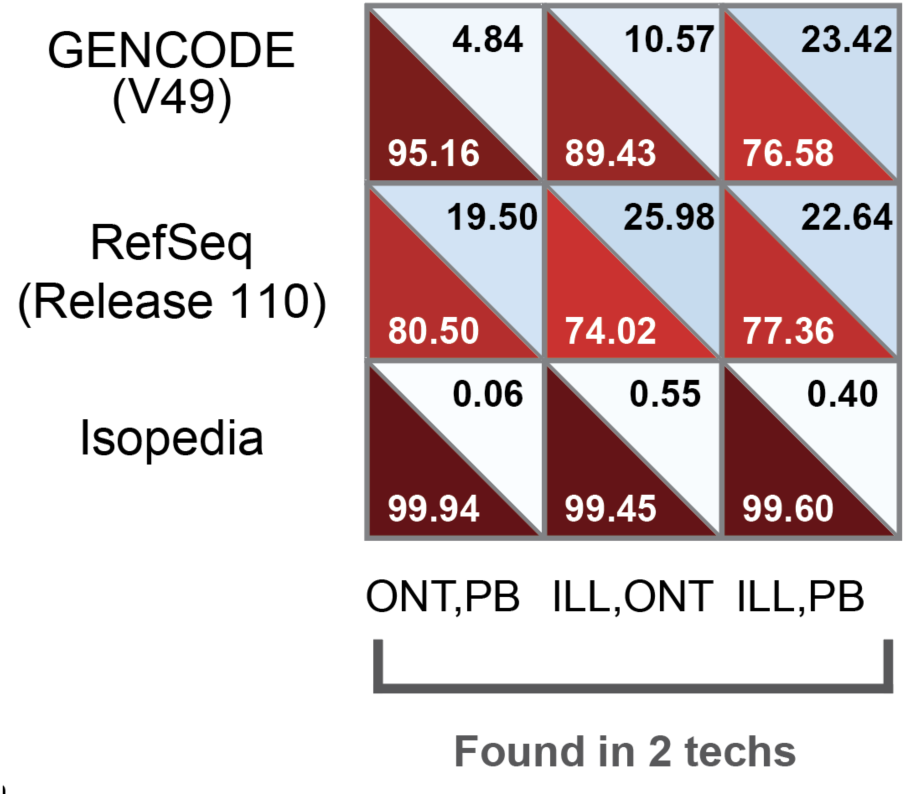
Classification of HG002 transcripts by annotation status using GENCODE V49 (first row), RefSeq Release 110 (second row), and Isopedia (third row). In each cell, the upper triangle shows the percentage of transcripts classified as novel, and the lower triangle shows the percentage classified as known. All transcripts identified by two technologies.

**Supplementary Figure S11.**
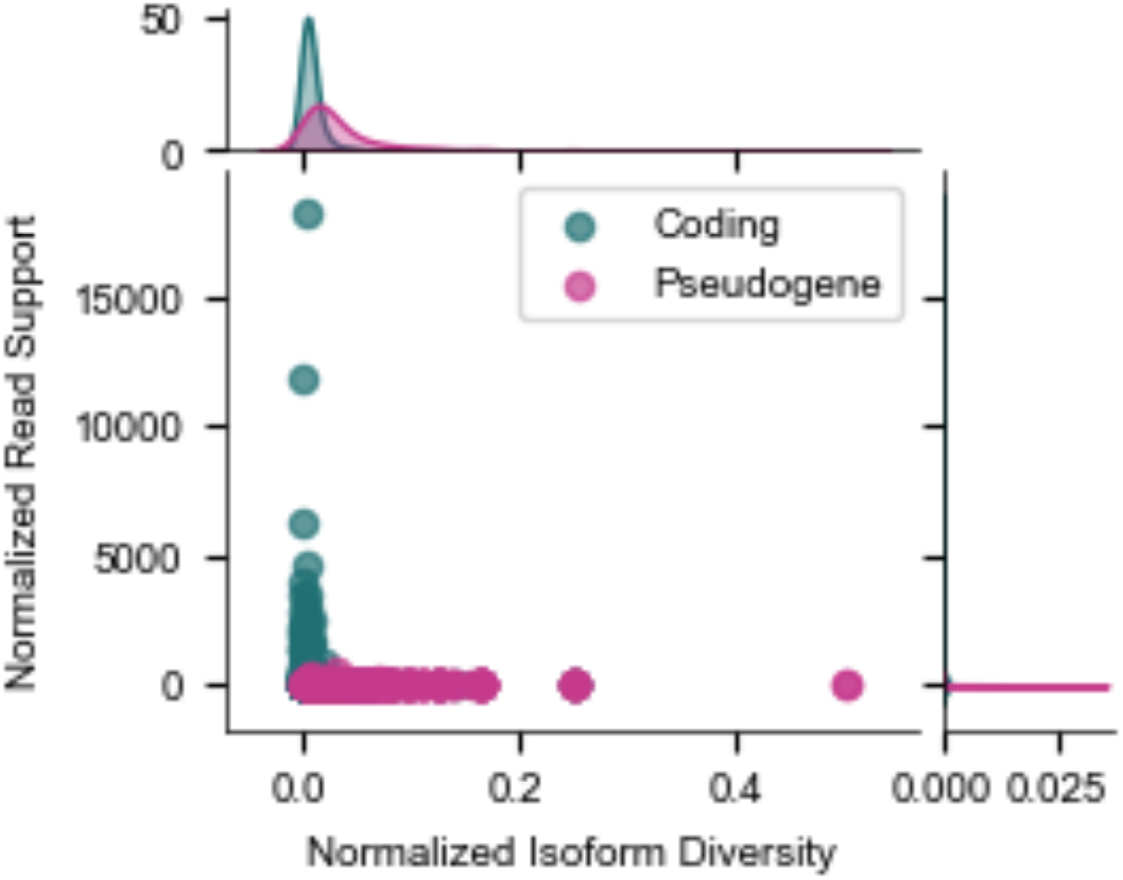
Normalized isoform diversity and expression of the pseudogene and its paired protein-coding gene. Top and right density plots summarize the distributions of NID and NSR, respectively.

**Supplementary Figure S12.**
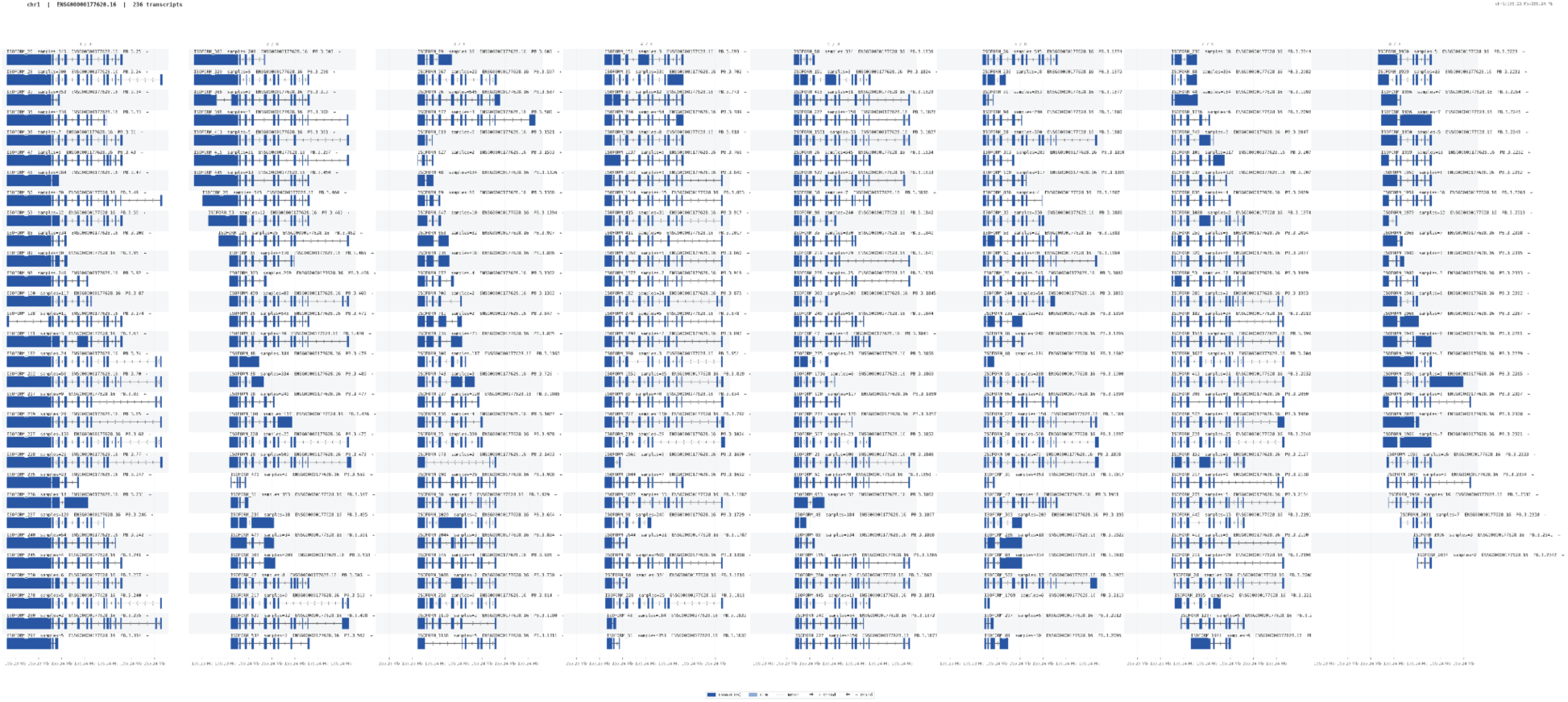
GBA1 transcripts that are validated by the isopedia catalog. The transcript was ordered by its coordinate of the genome. The label format is isoform id, sample size, annotated gene id, transcript id and strand.

**Supplementary Figure S13.**
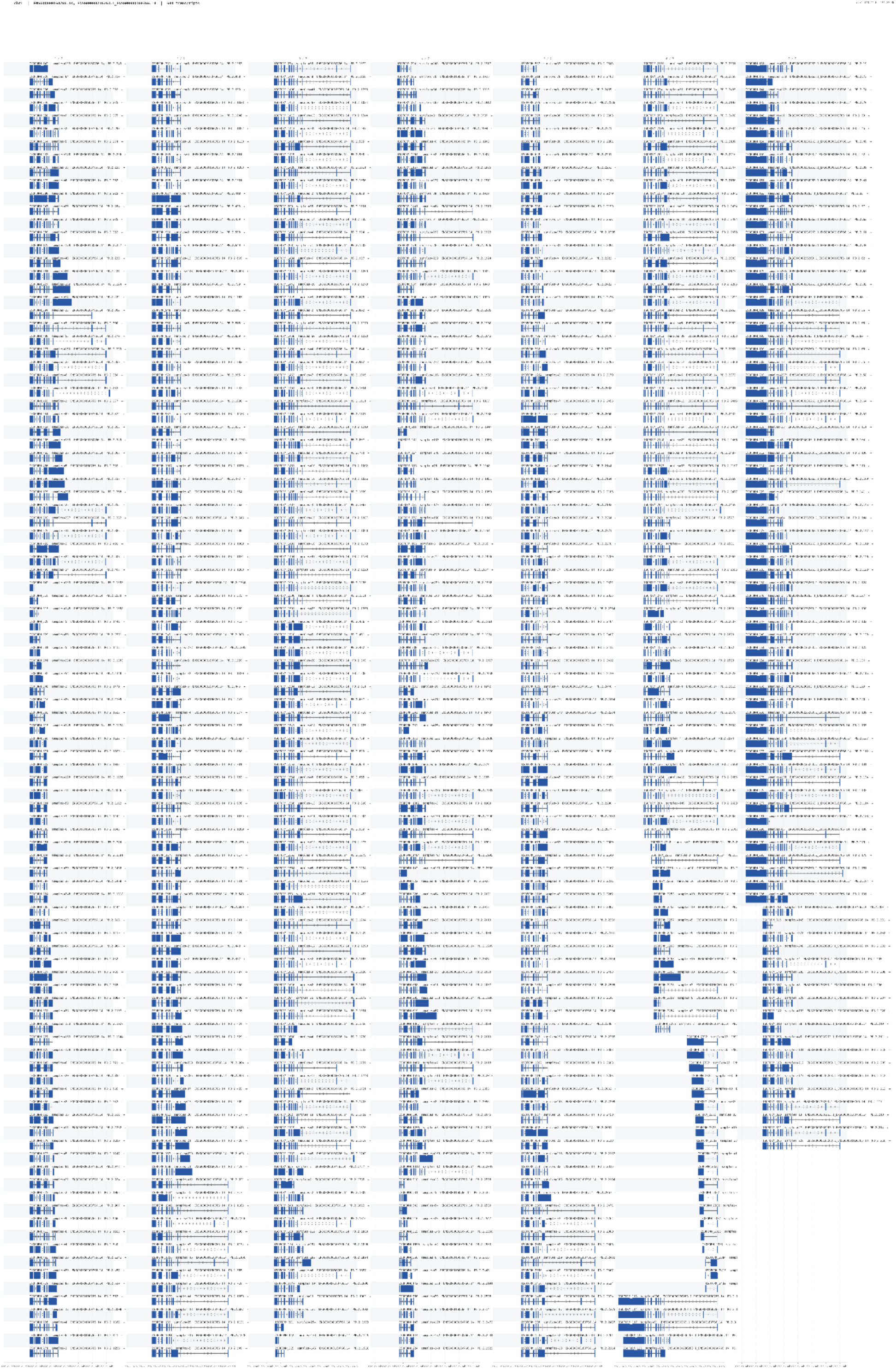
GBAP1 transcripts that are validated by the isopedia catalog. The transcript was ordered by its coordinate of the genome. The label format is isoform id, sample size, annotated gene id, transcript id and strand.

## Supplementary Note

### Note 1: Benchmark of Isopedia using LRGASP real datasets

We utilized the real sequencing datasets from LRGASP^1^ to evaluate Isopedia performance. The real sequencing datasets were derived from the WTC11 cell line and comprise six combinations of library preparation methods and sequencing technologies, each with three replicates. SIRV-Set4 transcripts were spiked into all datasets. For the simulated datasets, the ground truth consists of a subset of transcripts from the GENCODE v38 annotation along with their corresponding coverage values.

We assessed the re-identification robustness of Isopedia across different library preparation methods and sequencing technologies in real world scenarios. Isopedia created an index using a total of 18 datasets from the WTC11 cell line, representing six combinations of four library preparation methods and two sequencing technologies, with three replicates for each combination. The same GENCODE v38 GTF file was used as input for Isopedia. Overall, there are 237,012 transcripts in the GENCODE v38 annotation, and we observed an average consistency rate(not detected in any of replicates or detected in all replicates) of 91.5% (defined as whether a transcript was detected across all three replicates) across the six categories. Among them, the PacBio-CapTrap combination achieved the highest consistency rate of 93.4%, while the ONT-cDNA combination showed the lowest rate at 89.9%. The standard deviation of the consistency rates across all six categories was 1.28%, indicating that Isopedia performs consistently across different library preparation methods and sequencing technologies (**Supplementary Fig. S8, Supplementary Table S3**). Notably, the detected isoforms are positively correlated with the total reads number in the datasets, which suggests the larger read depth benefits the re-identification performance(**Supplementary Fig. S2**). The 18 WTC11 datasets were again used for assessing the quantification robustness within replicates from different library types. The mean coefficient variation(CV) was used to evaluate the quantification performance. We obtained an average of 0.21 CV among all library types, ranging from 0.181(PacBio-CapTrap) to 0.261(ONT-cDNA)(**Supplementary Fig. S3**). We evaluated Isopedia’s isoform re-identification performance on real datasets using SIRV transcripts and observed near-complete recall across all library types and replicates, with all spliced transcripts consistently recovered and only minor losses for a small number of unspliced transcripts in a few samples (**Supplementary Fig. S6**, **Supplementary Table 11**). We compared Isopedia’s consistency against other state-of-the-art tools across six LRGASP samples. IsoQuant achieved a mean consistency of 0.919 (0.883-0.945) and a mean CV of 0.145 (0.098-0.210). Oarfish achieved a mean consistency of 0.915 (0.890-0.927) and a mean CV of 0.159 (0.133-0.210). StringTie achieved a mean consistency of 0.861 (0.818-0.896) and a mean CV of 0.252 (0.186-0.339).

### Note 2: *GBA1/GBAP1 analysis*

We further explored the Isopedia-reidentified transcripts from Gustavsson et al.’s candidates by examining the sample count and expression level, measured as the average read count per sample. As shown in **Supplementary Table S13**, **Supplementary Table S14**, a substantial fraction of *GBA1* and *GBAP1* transcripts were supported by more than one read per sample on average. To assess the biological validity of the identified transcripts in the GBA1/GBAP1 locus, we applied three filters to the original transcript set: (1) removal of mono-exonic transcripts, (2) removal of transcripts mapped to the wrong strand, and (3) a minimum sample recurrence threshold. With a minimum sample requirement of 5, we retained 58 isoforms (193 transcripts) for *GBA1* and 199 isoforms (520 transcripts) for *GBAP1*. To further evaluate the functional potential of the *GBA1* transcripts, we applied ORFanage^2^ to annotate putative CDS regions, finding that 188 of 193 transcripts (97.4%) harbored a CDS, suggesting the vast majority have protein-coding potential. When relaxing the sample threshold to 2, the numbers increased to 86 isoforms (236 transcripts) for *GBA1* (**Supplementary Fig. S13**) and 303 isoforms (684 transcripts) for *GBAP1* (**Supplementary Fig. S14**), with ORFanage identifying CDS regions in 228 of 236 *GBA1* transcripts (96.6%). The consistently high proportion of CDS-containing transcripts across both thresholds supports the biological relevance of the retained *GBA1* isoforms, while the greater isoform diversity observed in *GBAP1* is consistent with its status as a transcribed pseudogene lacking protein-coding constraint.

We then leveraged the Isopedia splice-query functionality to capture additional isoform diversity by identifying transcripts containing at least one splice junction overlapping annotated GENCODE *GBA1/GBAP1* junctions. This approach identified 146 and 356 isoforms for GBA1 and *GBAP1*, respectively. To summarize isoform architecture in a compact and interpretable manner, we constructed splice-junction graphs for *GBA1* and *GBAP1* (**Supplementary Fig. S5**). We further quantified isoform complexity using an entropy-based metric (see **Supplementary Methods**). *GBAP1* exhibited higher isoform complexity (entropy = 0.68) than *GBA1* (entropy = 0.59), consistent with reduced protein-coding constraint at the pseudogene locus.

## Supplementary Methods

### Entropy measurement of *GBA1*/*GBAP1* splice graph

We defined an entropy-based measurement to evaluate the complexity of isoforms from GBA1 and *GBAP1*:

the weight is defined as:

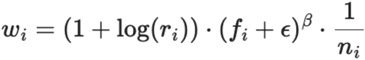

Where *r_i_* is the number of reads supporting isoform *i*, *f_i_* is the fraction of sample support isoform *i*, *n_i_* is the number of splice junctions in isoform *i*.

probability of isoform *i* is defined as:

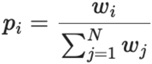

Entropy of the splice junction graph is defined as:

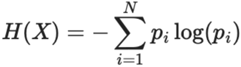

Normalized entropy is defined as:

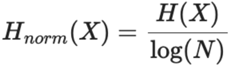

